# Distinct principles of genome compartmentalization in *Drosophila* and humans revealed by osmotic stress

**DOI:** 10.64898/2026.04.02.716189

**Authors:** Bright Amankwaa, Christopher Playter, Emily Stow, Jacob T. Sanders, Tianchun Xue, Rachel Patton McCord, Mariano Labrador

## Abstract

The three-dimensional organization of eukaryotic genomes into compartments, topologically associating domains, and loops is mediated by architectural proteins whose organizational principles vary across species. In *Drosophila*, insulator proteins including Su(Hw) and the histone variant γH2Av form liquid-liquid phase separation (LLPS) condensates, yet how this phase separation capacity relates to genome compartmentalization has remained unclear. Here we use hyperosmotic stress to simultaneously displace architectural proteins from chromatin in both *Drosophila* and human cells, enabling a comparative dissection of genome organization principles across species. We find that although human CTCF shares predicted LLPS properties with *Drosophila* insulator proteins, it does not form condensates upon osmotic stress, while *Drosophila* insulator proteins do. Hi-C analysis reveals that osmotic stress causes loss of compartments, TAD boundary strength, and loops in both organisms, but genome recovery after stress is rapid and near-complete in human cells while remaining substantially incomplete in *Drosophila* after one hour. Analysis of this recovery asymmetry reveals a fundamental difference in compartment organization: whereas human A and B compartments engage in robust homotypic long-range interactions, *Drosophila* B compartments rarely participate in long-range B-to-B contacts, indicating that the *Drosophila* genome does not replicate canonical A/B compartment organization. Instead, *Drosophila* genome architecture is dominated by A-to-A interactions, and A compartments are specifically enriched in γH2Av and Su(Hw), with moderate enrichment of cohesin subunits. Furthermore, loops in *Drosophila* are mechanistically independent from compartments and TADs, recovering before compartment structure is restored, and are anchored by Su(Hw) and cohesin rather than by dCTCF. Together, these findings suggest that the LLPS properties of γH2Av and Su(Hw) underlie A compartment formation in *Drosophila* through a mechanism distinct from the heterochromatin-driven B compartment interactions that predominate in vertebrates, revealing fundamentally different organizational principles between fly and human genomes.

## Introduction

The three-dimensional (3D) genome organization of eukaryotes serves as a crucial epigenetic regulatory landscape and complements the encoded information in the linear genomic sequence. Aberrations in this spatio-temporal organization of the genome are implicated in a number of developmental disorders and diseases (Ahn et al., 2021; Lupiáñez et al., 2015). Therefore, understanding the mechanisms mediating the establishment and maintenance of the 3D genome organization is critical to unravelling fundamental mechanisms of gene transcription, replication, and DNA damage repair. At least three length scales of the eukaryote chromosome folds have been characterized (Lieberman-Aiden et al., 2009; Mirny et al., 2019). First is the preferential segregation of chromosomes into chromosomal territories. Second is the demarcation of the genome into global active (A) and inactive (B) compartments. Finally, at the local scale, folds form interacting domains referred to as topologically associating domains (TADs) (Kentepozidou et al., 2020).

In addition to histone modifications, architectural proteins such as insulator binding proteins (IBPs), cohesin, and specific transcription factors play a key role in orchestrating the assembly of the three-dimensional genome by defining domain boundaries and stabilizing the folded chromatin loops (Misteli, 2020; Rao et al., 2014). One of the key genome organizing functions of IBPs is their control of the effect of regulatory elements, such as enhancers, on gene promoters (Ali et al., 2016; Furlong and Levine, 2018). Most insulator proteins such as Su(Hw), CP190, Mod(mdg4)67.2, BEAF-32, and the *Drosophila* CCTC-binding factor (dCTCF) have been identified in *Drosophila* (Matthews and White, 2019). In contrast, among mammals, CTCF is the only well characterized insulator protein (Matthews and White, 2019).

Two major mechanisms have been proposed to explain the principle through which architectural proteins modulate the spatio-temporal organization of the genome (Mirny et al., 2019; Nuebler et al., 2018). First is the passive segregation of chromatin into distinct compartments based on shared epigenetic characteristics (Falk et al., 2019). The second mechanism relates to the active interplay between cohesin and the CCCTC-binding factor (CTCF) via what is referred to as the loop extrusion process (Fudenberg et al., 2016). Several lines of evidence suggest that liquid-liquid phase separation (LLPS) plays a role in the segregation of shared epigenetic features into compartments (Alberti et al., 2019). Previous reports suggested that compartments emanate hierarchically from TADs (Lieberman-Aiden et al., 2009; Rao et al., 2014). However, recent observations demonstrate that, at least in mammals, they are mechanistically decoupled structures (Mirny et al., 2019; Nuebler et al., 2018). In fact, the process of loop extrusion, which helps form TADs, counteracts compartmentalization, and removing TAD mediating factors increases compartment formation (Nuebler et al., 2018; Rao et al., 2017; Schwarzer et al., 2017).

Whereas 3D genome features appear to be conserved across species, notable distinctions exist within and between organisms (Acemel and Lupiáñez, 2023). For instance, CTCF-cohesin loops appear to only represent a subset of domains in mammals and the non-CTCF-cohesin mediated domains do not show the classical loop extrusion features (Szabo et al., 2020). Moreover, in *Drosophila*, domains with loop extrusion features are infrequent (Messina et al., 2023). In addition, the CTCF N-terminal region required to interact with the cohesin PDS5A domain and therefore block loop extrusion is not conserved in *Drosophila* (Acemel and Lupiáñez, 2023; Nora et al., 2020).

These findings imply that the loop extrusion process could be more significant in organizing the vertebrate genome, suggesting further that alternative mechanisms may play a more prominent role in shaping the *Drosophila* genome. We have shown that in response to hyperosmotic stress, *Drosophila* insulator proteins demix with cohesin subunits forming condensates that exhibit LLPS properties, including susceptibility to the LLPS disrupting agent, 1,6-hexanediol (Amankwaa et al., 2022). Although human CTCF (hCTCF) also undergoes a form of *in vivo* clustering, LLPS appears to play only a modest role (Shi et al., 2021). For instance, hCTCF shows only marginal susceptibility to the LLPS disrupting agent, 1,6-hexanediol (Lee et al., 2022). The response of the human and *Drosophila* whole genomes and their architectural factors to osmotic stress could therefore serve as an important tool to understand general genome organization principles.

It remains unclear whether human CTCF exhibits a response to salt stress, leading to the formation of CTCF condensates, akin to the observed phenomenon in *Drosophila*. Moreover, whereas it has been shown that hyperosmotic stress induces a reversible loss of genome structure in humans (Amat et al., 2019), its effect on the *Drosophila* genome structure is less understood (Irianto et al., 2013; Khatibi et al., 2019; Schoborg et al., 2013). We aimed to harness the osmotic stress response to gain insights into genome organization principles across several scales in both human and *Drosophila*. Our work sought to specifically assess the hyperosmotic stress effect on the 3D genome landscape with regards to the self-interacting domains, compartments, and loop structures in both *Drosophila* and human cells. Considering the observed variations in the role of cohesin-CTCF mediated genome folding in mammals and *Drosophila*, we anticipated distinct patterns of genome restructuring in response to osmotic stress within these two genomes.

We found that although hCTCF and *Drosophila* insulator proteins share comparable predictors of LLPS and ensemble properties, hCTCF does not form osmotic stress induced bodies akin to those formed by *Drosophila* insulator proteins. However, following osmotic stress, both cell types undergo chromatin shrinkage and an overall maintenance of chromosome territories. Also, in both cell types, we observed loss of compartments and the weakening of TAD boundary strength throughout the genome as well as a significant increase in non-compartmental interactions in the long-range.

Interestingly, whereas the genome organization in human cells almost fully recovers after 1 hour in normal media post osmotic stress, the recovery of *Drosophila* cells after 1 hour in normal media is incomplete as genomes fail to regain normal compartment structure and TAD boundary strength remains significantly reduced. Analysis of these differences revealed that the *Drosophila* genome does not replicate the canonical A/B compartment organization found in vertebrates. In *Drosophila*, A compartments engage in A-to-A long-range interactions in a manner similar to that found in the human genome. However, B compartments fail to reach long-range B-to-B interactions. Interestingly, A compartments are specifically enriched in γH2Av and Su(Hw) insulator proteins, suggesting that the LLPS properties of these proteins may be required for compartment formation. We also found that loops are independent from TADs and compartments in *Drosophila* since recovery of loops is not accompanied by simultaneous recovery of compartments or TAD boundary strength.

## Results

### Human and *Drosophila* genomes display similar osmotic stress-induced chromatin shrinkage events but reveal differences in architectural protein responses

We first compared the global response of the human genome and *Drosophila* genome to hyperosmotic stress. For both human A375 and *Drosophila* S2 cells, we changed from standard media (“control”) to media containing 250mM NaCl and incubated for 1 hour (“stress”). We then replaced the high salt medium with standard medium for another 1 hour (“recovery”). When the cells were fixed and stained with DAPI and CTCF antibodies, we observed that osmotic stress significantly reduced chromatin volumes in both cell types (Figure 1B). The human cells regained their original chromatin volume when allowed to recover in the physiological media (Figure 1B). In contrast, there was a significant difference in the DAPI area between the control and recovery for S2 cells (Figure 1D). This suggests that the mechanisms involved in reconstructing genome structure or the physiological conditions leading to full recovery may differ between flies and humans.

**Figure 1:**
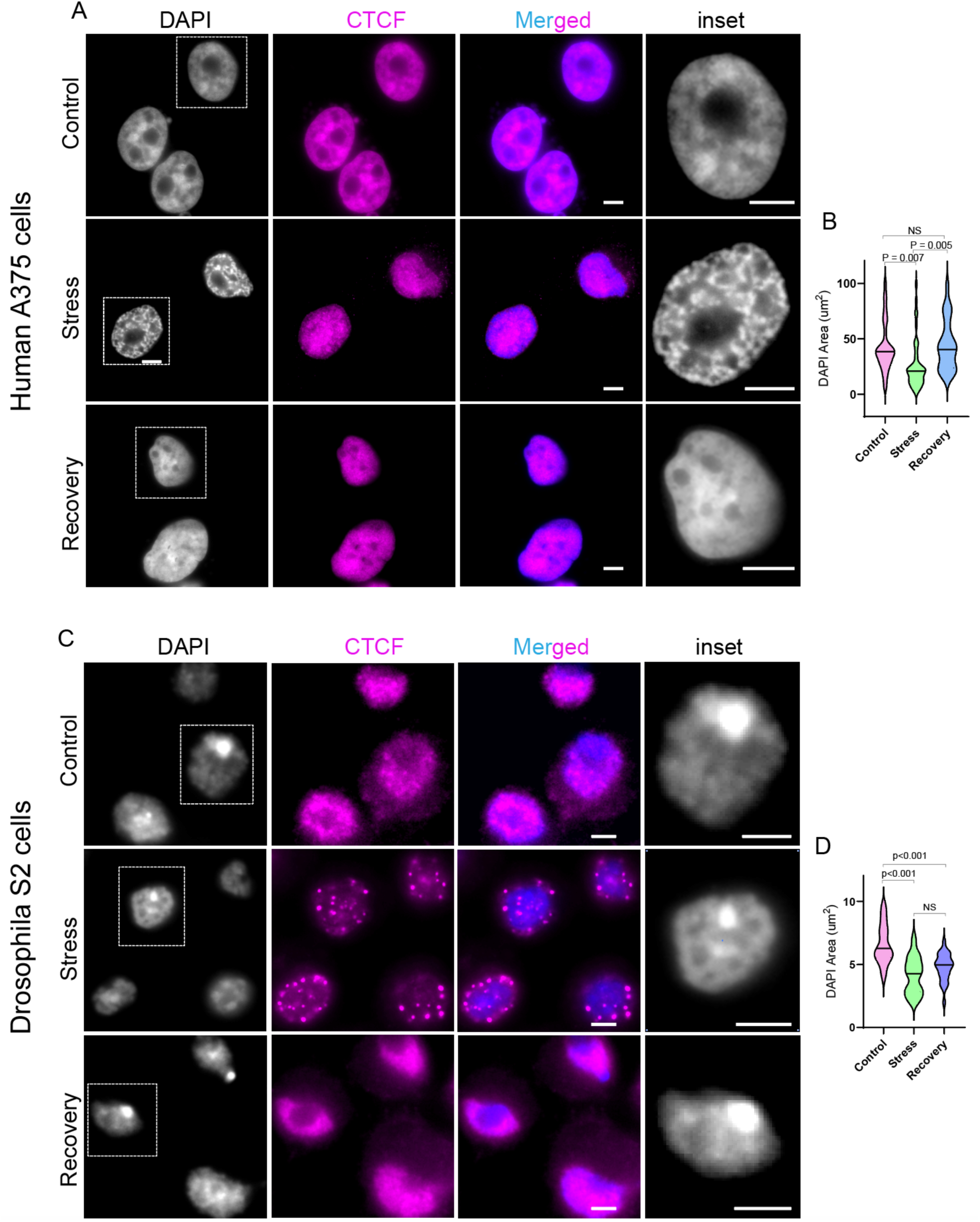
Distribution of architectural proteins reveals differences in the response to osmotic stress between *Drosophila* and human genomes. **A.** Human A375 cells immuno-stained with CTCF (magenta) and counterstained using DAPI. In control cells (top panel), grown in normal media, CTCF appears uniformly distributed on chromatin. In stress cells (middle panel), grown in media containing 250mM NaCl for 1 hour, chromatin appears highly condensed (inset) and occupancy of CTCF on chromatin is reduced, as it migrates to the nucleolus, interchromosomal spaces and the nuclear periphery. In these cells, CTCF does not form visible condensates. Recovery cells (bottom panel) were first grown in media containing 250mM NaCl for 1 hour and then returned to physiological media for 1 hour. Recovery cells look similar to control cells as CTCF is uniformly distributed on the chromatin and chromatin condensation appears similar to control (inset). **B.** Comparison of chromatin volume (DAPI stained area) for media (control, N = 30), 250mM NaCl treated (stress, N = 30) and physiological media treatment after 250mM NaCl treatment (recovery, N=30). Significant difference (One-Way ANOVA followed by multiple comparisons) in chromatin volume exists between control and stress (p = 0.007), stress and recovery (p = 0.005) but not between control and recovery. **C.** *Drosophila* S2 cells are uniformly immuno-stained with dCTCF (magenta) in normal media (control, top panel). In response to 250mM NaCl treatment (stress), chromatin appears condensed (DAPI, inset). The occupancy of dCTCF on the chromatin is reduced as it forms insulator bodies and migrates to nucleoplasm spaces. Salt stress treated cells returned to physiological media (recovery) look similar to the control cells as dCTCF is uniformly distributed on the chromatin. Scale bar = 2µm **D**. Comparison of chromatin volume (DAPI stained area) for media (control, N= 30), 250mM NaCl treated (stress, N= 30) and media treatment after 250mM NaCl treatment (recovery, N= 30). A significant difference exists in chromatin volume between control and stress (p <0.001), control and recovery (p <0.001) but not between stress and recovery. Scale bars = 2 µm.

Considering that hyperosmotic stress induces demixing of *Drosophila* IBPs forming condensates with LLPS features (Amankwaa et al., 2022; Schoborg et al., 2013), we wondered whether hCTCF would also respond to NaCl-induced stress in a similar fashion. Our analysis showed that contrary to the *Drosophila* insulator proteins, treating human A375 cells with NaCl for 1 hour failed to induce hCTCF bodies (Figure 1A and 1C). This is surprising since hCTCF displays similar residue charge distribution patterns, disorder tendencies, and predicted ensemble properties as those found in *Drosophila* IBPs (Supplemental Figure S1).

In summary, osmotic stress causes significant chromatin volume reduction in both species accompanied by condensate formation of Drosophila IBPs, but not human CTCF. Following return to normal media, human cells fully restore pre-stress chromatin volume within one hour, while *Drosophila* cells genome only undergo a partial expansion.

### Osmotic stress induces distinct global changes in DNA contacts in *Drosophila* and human chromosomes

To compare genome-wide chromosome contacts, we performed Hi-C in A375 and S2 cells, under control, stress, and recovery conditions (Arima protocol, see Methods and Supplemental Table 1). At 1 Mb resolution, chromosome territories were largely maintained in both species following osmotic stress (Figure 2A). Despite this, osmotic stress caused a consistent shift in contact frequencies: short-range interactions where lost while long-range interactions were gained in both cell types, confirmed by log2 (Stress/Control) ratio heatmaps (Figure 2B, and 2C). In *Drosophila* S2 cells, this manifested as a dramatic loss of intra-centromeric and gain of centromeric interactions through the chromosome arms (Figure 2C).

**Figure 2.**
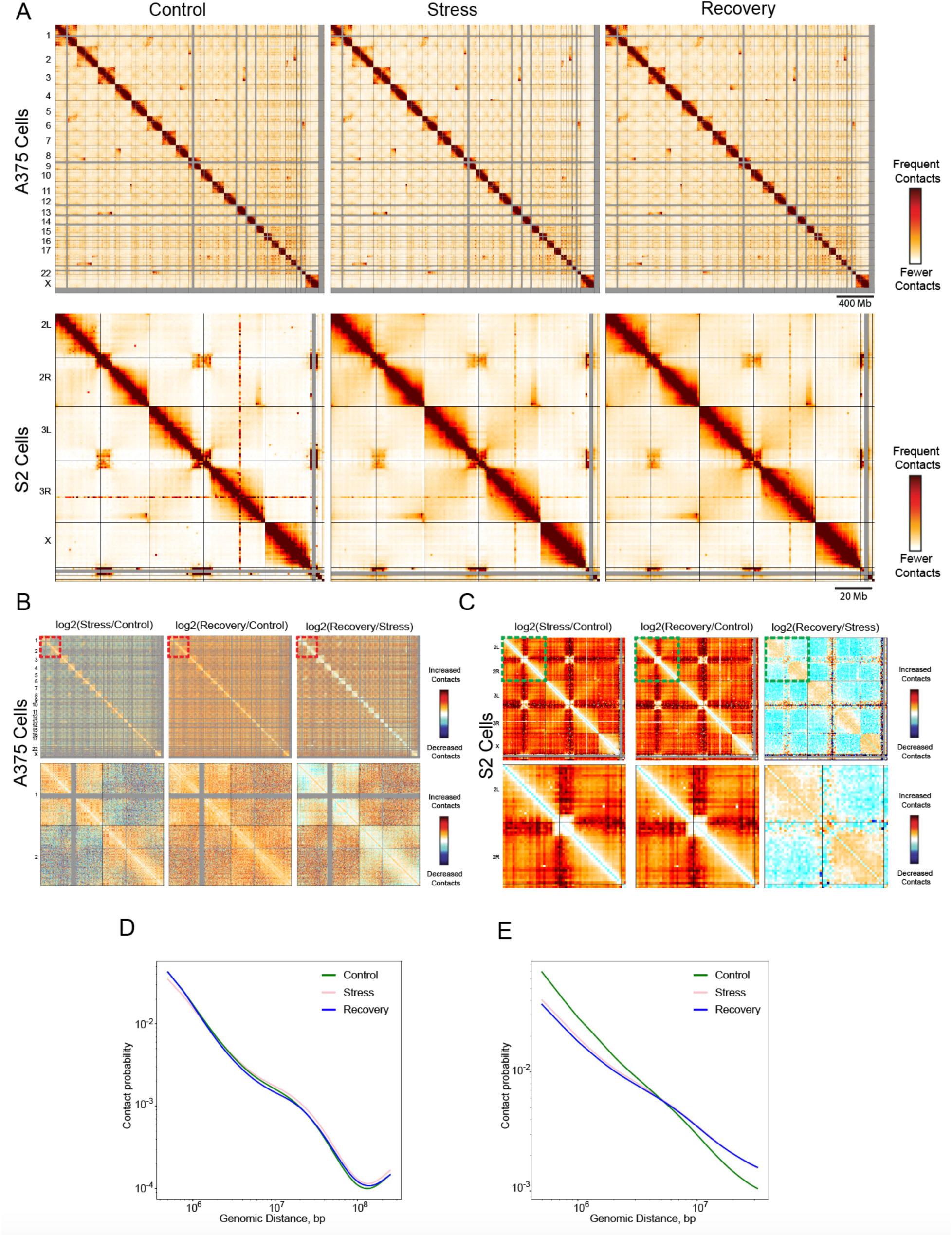
Hi-C reveals differences in osmotic stress response between *Drosophila* and human genomes. **A.** A 1Mb binned heatmap of human A375 cells (top panel) and *Drosophila* S2 cells (bottom panel) across the 3 conditions. **B**. 1.0 Mb resolution log2ratio whole genome heatmaps of human A375 Stress versus Control (left panel), Recovery versus Control (middle panel), and Recovery versus Stress (right panel). Insets of chromosomes 1 and 2 are placed below their respective heatmaps. **C.** 1.0 Mb log2ratio whole genome heatmaps of *Drosophila* S2 Stress versus Control (left panel), Recovery versus Control (middle panel), and Recovery versus Control (right panel). Insets of chromosome 2L and 2R are placed below their respective heatmaps. **D.** A scaling plot showing contact decay with distance across all chromosomes at a 250 kb bin size in human A375 cells. **E**. A scaling plot showing contact decay with distance across all chromosomes at a 1Mb bin size in *Drosophila* cells

The two species differed in their inter- versus intra-chromosomal stress responses (Figure 2D and 2E). In human A375 cells, chromosome territories condensed and moved apart, increasing intra-chromosomal interactions and decreasing inter-chromosomal interactions. In *Drosophila* S2 cells, distal interactions increased both within chromosomes and between chromosomes. Contact frequency decay curves confirmed these patterns: both species showed loss of sub-1 Mb contacts, which is consistent with the results of polymer models simulating the loss of loop extrusion factors from chromatin (Fudenberg et al., 2017), but their recovery trajectories diverged sharply. Human cells recovered to near control levels, with long -range interactions (10 Mb) falling below the control during recovery (Figure 2D) and only very long-range contacts (100 Mb) remaining slightly elevated, which matches a persistent elevation in contacts between chromosomes. *Drosophila* cells, in contrast, showed no meaningful recovery as the stress and recovery decay curves were nearly identical, with persistent loss of local contacts and elevated distal interactions (Figure 2E).

In human A375 cells, the stress response was gene-density dependent: smaller, gene-dense chromosomes (from chr14 to chr22) gained more *trans*-interactions than larger gene-poor chromosomes (Figure 2B). Chr18, which is a similar size but gene-poor did not gain interchromosomal interactions, confirming this is a gene-density rather than size effect. Strikingly, this pattern is similar to the global *Drosophila* response, suggesting that gene density may be a conserved determinant of the genome’s osmotic stress response across species.

Overall, these results indicate that osmotic stress causes a conserved shift from local to distal contacts in both species, but that genome recovery is far more complete in human than *Drosophila* cells, and that the balance of *cis* versus *trans* contact changes differs substantially between species.

### Hyperosmotic stress disturbs the A/B compartment organization in human and *Drosophila* genomes with varying degrees of recovery after stress

Considering the variations observed in the global rearrangement of genome contacts following osmotic stress, we wondered whether similar remodeling variations exist between *Drosophila* and human cells at higher resolution, such as at the level of compartments.

It was evident from the contact probability maps that in both cell types osmotic stress results in loss of the characteristic long-range compartment interactions, including a loss in the human genome’s alternating ‘plaid-like’ compartment pattern (Figure 3A). Contrary to the well-defined compartment borders in the control cells, hyperosmotic stressed genomes were marked by less distinct and blurry borders (Figure 3A). This indicates higher levels of interactions between compartment types after osmotic stress. Interestingly, compartments appeared mostly restored in the A375 recovery sample, as observed previously in a less strong osmotic stress (100 mM salt) in a different human cell type (Amat et al., 2019), but the restoration was less complete in the *Drosophila* S2 cells (Figure 3A and 3B).

**Figure 3.**
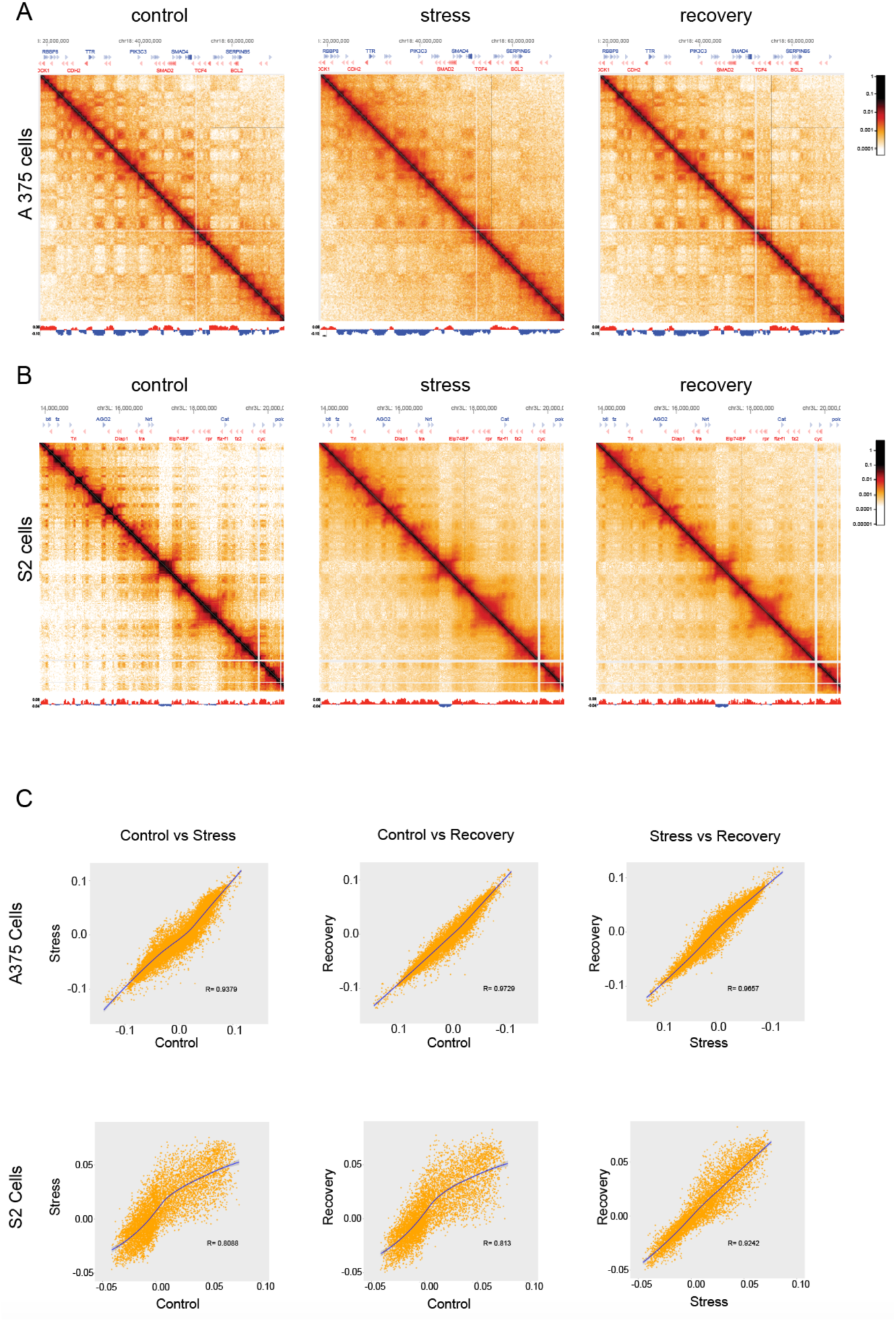
*Drosophila* and human display differences in the degree of A/B compartment readjustment after perturbation with hyperosmotic stress. **A.** Heatmaps of genomic contacts (250kb resolution) of chromosome 18: 20Mb-70Mb of human A375 media treated, 250mM NaCl treated and recovery. The first principal component, aligned below each matrix, shows compartment classification into A (red) and B (blue) compartments. **B.** Heatmaps of genomic contacts (20 kb resolution) of chromosome 2L:14Mb-20Mb of *Drosophila* S2 media treated, 250mM NaCl treated and recovery are shown. The first principal component, showing compartment identity, is aligned below each heatmap as in (A). **C.** Genome-wide Pearson’s correlation of first principal component values for human (top panel) or *Drosophila* (bottom panel) comparing control versus stress (far left panel), control versus recovery (middle panel), and stress versus recovery (far right panel). The lines of best-fit are shown in blue through the data points. The blue lines through the data points indicate the lines of best fit.

We next classified genomic regions into A or B compartments using principal components analysis (PCA) on the 250 kb (human) or 20 kb (*Drosophila*) Hi-C matrices. In this context, positive PC1 values signify A compartments, typically associated with gene-rich regions (euchromatin), while negative values indicate B compartments associated with gene poor regions (heterochromatin). Despite the visible disruption of the compartment patterns, we found a strong correlation in the compartment identity across all three conditions in both organisms (Figure 3C). This suggests that while hyperosmotic stress alters compartment integrity, these modifications are not sufficiently drastic to result in their complete rearrangement. In general, however, across the various conditions, the PC1 values correlated more strongly in the A375 cells than in the S2 cells (Figure 3C). We infer from this that compartment segregations in the *Drosophila* genome are more susceptible to osmotic stress compared to those in the human genome. In both species, osmotic stress caused the largest decrease in compartment correlation (r = 0.809 for S2, r = 0.923 for A375). In S2 cells, the correlation was strongest in stress vs. recovery (r = 0.9242) while it A375 cells the control vs. recovery conditions were the most correlated (r = 0.973) (Figure3C). These results also suggest that compartment reassembly after osmotic stress is either more robust or faster in the human A375 cells compared to the *Drosophila* S2 cells.

### Human and *Drosophila* genomes exhibit distinct compartment identity and compartment strength plasticity in response to osmotic stress

While the correlation results above show that compartment identity is largely conserved during osmotic stress, minor switches in compartment identity can have biological implications and associations with changes in gene expression (Dixon et al., 2012; Li et al., 2024). Several studies have emphasized the impact of such minor switches and their consequences under various stressors like heat and salt (Li et al., 2024). We wondered whether the decreased compartment segregation triggered by osmotic stress would cause compartment identity changes in the S2 and A375 cell genomes.

We performed a genome wide comparison of compartments across conditions, classifying a bin as “switching compartments” if the PC1 sign changed from negative to positive or vice versa and the value shifted at least 10% of the full PC1 range. We found that A375 cells exhibited a lower percentage of B to A transitions (0.94% of genomic bins) and a higher percentage of A to B transitions (4.75%) during salt stress (Figure 4A, left, Supplemental Figure S2A). In contrast, S2 cells showed more B to A transitions (10%) than A to B (1.15%) during this stress condition (Figure 4B, left,, Supplemental Figure S2B). Another striking difference between human and *Drosophila* cell compartment changes was the degree to which these switches recover after osmotic stress. We classified switches as “recovered” if they were no more than 2% different than their Control PC1 value in the Recovery condition and “not recovered” if they remained within 2% of their Stress value during the Recovery condition. In A375 cells, 18% of switched bins fully revert back to their original state and a further 71% partially recover, leaving only 11% of all switched bins unrecovered after the return to normal osmotic conditions (Figure 4A, middle and right panels). However, in S2 cells, only 3% of bins fully recover and 35% of switches do not recover at all from their osmotic stress compartment state (Figure 4B, middle and right panels).

**Figure 4.**
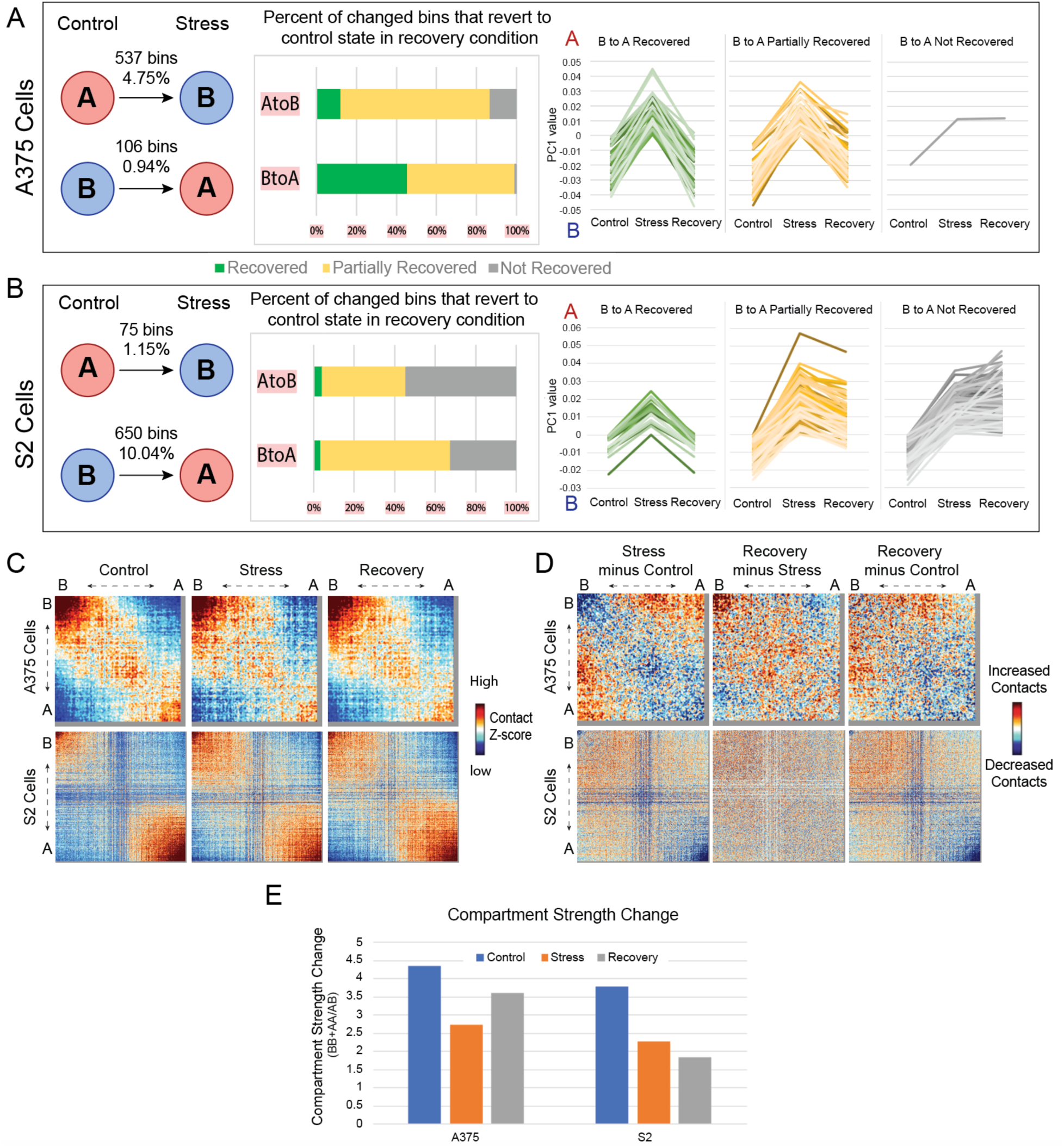
The human and *Drosophila* genomes differ in compartment strength and identity changes after osmotic stress. **A.** For A375 cells, the left panel shows the number of 250 kb bins and percent of the whole genome that switch compartments, as defined by a PC1 sign change and at least a 10% shift in PC1 value. Given these compartment switch bins, the middle panel shows the proportion that revert fully (green), partially (yellow), or not at all (grey) upon return to control conditions. “Recovered” is defined as a Recovery condition PC1 value within 2% of the Control value while “Not Recovered” bins have a Recovery PC1 value within 2% of the Stress value. The patterns of PC1 value change are shown in the graphs at right, where each line represents the trajectory of one genomic bin. **B.** Same analysis as in A but for *Drosophila* S2 cells **C.** Saddle plots generated from 250Kb and 20kb binned distance corrected Z-score matrices for chromosome 18 of human A375 cells (top panel) and chromosome 2L of *Drosophila* S2 cells (bottom panel) respectively. The binned matrices in each case are reordered from strongest B to strongest A compartment identity. The binned matrices were smoothed to 500Kb for the human A375 cells while the *Drosophila* S2 cell genome was smoothed at 20kb. **D.** Saddle plots showing comparison of A and B compartment interaction strengths in different conditions for human A375 cells (top) and *Drosophila* S2 cells (bottom). The Z-score interaction strengths were subtracted between the indicated conditions and the differences are shown. **E.** Compartment strength recovers in A375 much more than in S2 cells. The compartment strength for each condition in each cell type is calculated as (AA+BB)/AB. Where AA, for example, is the sum of all interactions within the top 20% of A compartment bins: the bottom right corner of the saddle plots.

To quantitatively assess changes in interaction frequency between or within compartments, we utilized saddle plot analysis to quantify the strength of homotypic (either A-A or B-B) or heterotypic (A-B or B-A) interactions. To generate the saddle plots, genomic distance corrected Z-score 20kb and 250Kb binned matrices for S2 and A375 cells respectively were reordered based on compartment strength (from strongest B to strongest A using eigenvector PC1 values as demonstrated elsewhere) (Sanders et al., 2022).

Reordering the compartment identities by their interaction strengths revealed an obvious separation between the regions of strong B and A contacts in both the A375 and S2 cells (Figure 4C). Strikingly, whereas the A375 control saddle plot shows comparable strengths of A-A and B-B homotypic interactions consistent with canonical compartment organization, the S2 control saddle plot reveals a marked asymmetry in which A-A interactions are substantially stronger than B-B interactions (Figure 4C). This asymmetry, already present under normal conditions, indicates that the *Drosophila* genome lacks the strong long-range B-to-B interactions that characterize vertebrate genome organization. As we saw in contact maps, we observed intermingling of compartments in the stressed A375 and S2 cells.

In A375 cells, we observed an overall loss of the strongest homotypic compartment interactions (A-A, and B-B) but a gain of heterotypic compartment interactions (A-B and B-A) after osmotic stress (Figure 4D, left panel). The most striking loss of interactions occurs within the B compartment. Upon comparing the recovery sample to the stress condition, we observed that after recovery, the previously lost homotypic B interactions were restored, while homotypic A interactions are still weak (Figure 4D middle panel). This results in a final state where B-B interactions are restored to the control state, A-A interactions are still weaker, and some A-B interactions remain increased after recovery as compared to the original control (Figure 4D right panel).

In contrast, in *Drosophila* S2 cells, the homotypic-A interactions are the most strikingly lost (Figure 4D, left panel). There is also loss of interactions between the strongest B compartments, but both weaker B-B interactions and heterotypic A-B compartment contacts increase following osmotic stress (Figure 4D). As with compartment switches, markedly less recovery of compartment strength is seen in S2 cells after return to original conditions. This is evident both in the negligible differences in the Stress and Recovery saddle plots (Figure 4D, middle panel) and when we quantified compartment strength in each condition as the sum of homotypic over heterotypic interactions (AA+BB/AB; Figure 4E).

### Hyperosmotic stress causes comparable major loss of TAD structures but varied recovery patterns between *Drosophila* and human genomes

To assess whether TADs are differentially affected by osmotic stress between *Drosophila* S2 cells and human A375 cells, we called TADs at 5kb and 40kb resolutions for the *Drosophila* S2 cells and human A375 cells, respectively.

Visual inspection of the Hi-C maps revealed the distinct blocks of triangular-shaped structures signifying individual TADs with intact boundaries in the controls of both species (Figure 5A). In the osmotic stress cells however, the TAD boundary integrity is compromised, resulting in the merging of neighboring domains into singular entities (Figure 5A and 5B). Notably, the disrupted TAD patterns under stress conditions appeared reestablished in the A375 recovery cells but not in the S2 cells (Figure 5A and 5B).

**Figure 5:**
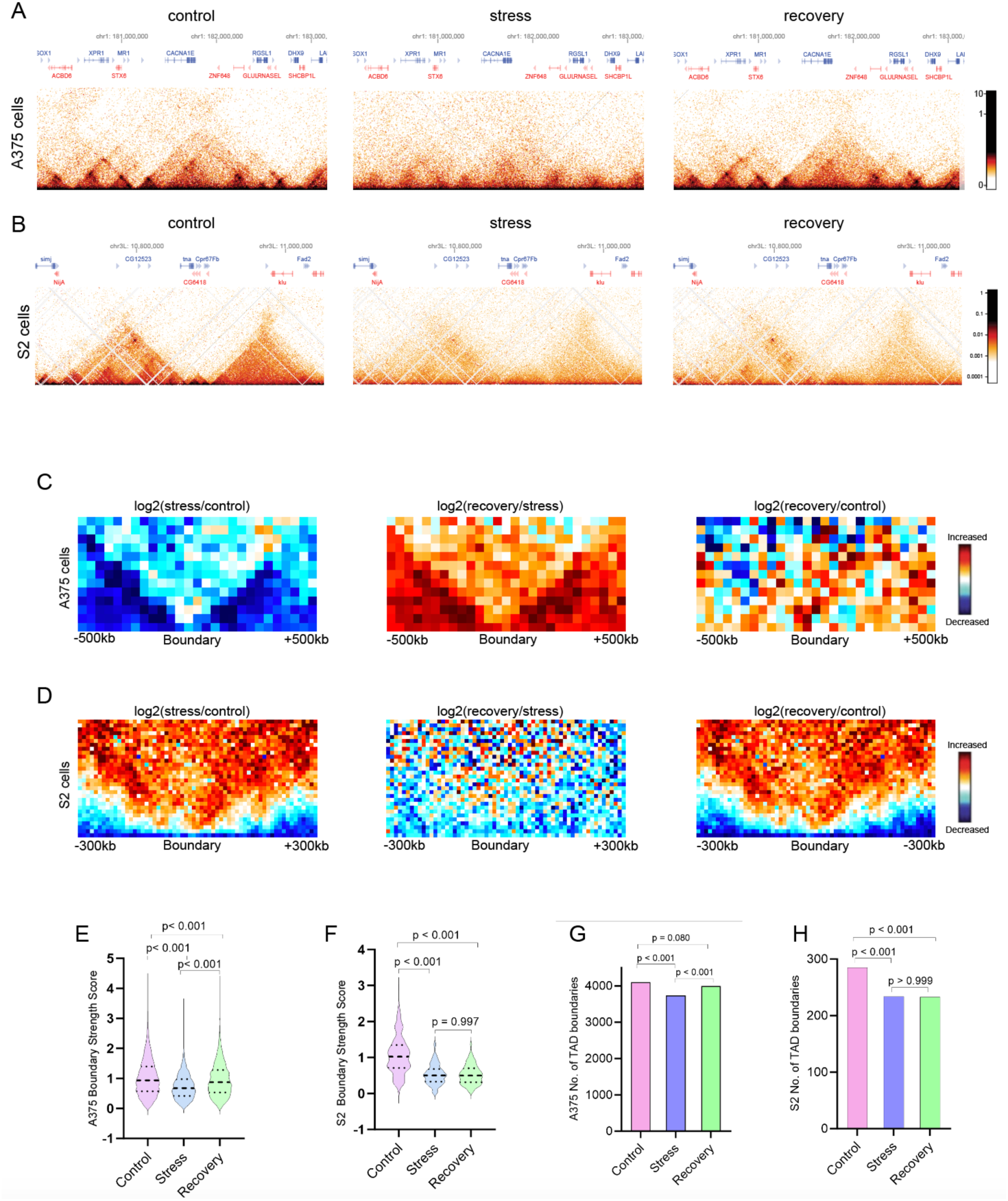
*Drosophila* and human genomes display distinct TAD changes in response to osmotic stress. **A.** Heatmaps showing human A375 TAD structures (called at 40kb) at chr1:180.2 - 183 Mb. **B.** Heatmaps showing *Drosophila* S2 cell TAD structures (called at 5kb) at chr3L:10.6 - 11 Mb. **C.** Log_2_Ratio of aggregate contact maps at TAD boundaries called at 40 kb bins in A375 cells. **D.** Log_2_Ratio of average TAD boundary (called at 5kb) plots from S2 cells. **E.** Human A375 cell comparison of TAD boundary strength scores. P-values less than 0.05 are deemed significant. **F.** Comparison of TAD boundary strength score in *Drosophila* S2 cells. P-values are generated by one-way ANOVA and values less than 0.05 are deemed significant. **C-F**. Boundary strengths were derived from the cworld-dekker method to calculate insulation scores. **G-H.** Comparison of numbers of detected TAD boundaries in human A375 cells (G) or *Drosophila* S2 cells (H) derived from the insulation scores. P-values were generated using Fisher’s exact test for each pair of conditions, with values less than 0.05 considered statistically significant.

The apparent merging of adjacent domains in the stressed condition is manifested in the significant reduction in number of TAD boundaries compared to the controls (Figure 5G, 5H, and Supplemental Figure S3). As observed with compartment changes, this TAD boundary count increases back to control levels in the A375 recovery condition but remains similar to the stress condition in S2 recovery cells.

To gain further insights into the integrity of the TADs in response to osmotic stress, we calculated the TAD separation score (insulation score) that allows for the assessment of the TAD boundary strengths, that is the degree of contact separation between TADs (Lajoie et al., 2015). We observed a significant reduction in boundary strengths in the stress compared to controls for both cell types (Figure 5E and 5F). However, whereas a significant difference existed between the human stress and recovery TAD boundary strengths (P <0.001) as the recovery cells become more similar to controls, we found no significant difference between the *Drosophila* stress and recovery TAD boundary strengths (P-value = 0.997) (Figure 5C and 5D). This reinforces the observed restoration of TAD structures in the A375 cells compared to those in the S2 cells. Nonetheless, the regained boundary strength following recovery in the A375 was quite not enough to bring it to the control level (Figure 5F).

To illustrate the changes in TAD separation, we plotted the average interaction profile around TAD boundaries (Figure 5C and 5D). In both organisms, our analysis revealed a notable decrease in contacts within TADs under osmotic stress conditions compared to the control, likely stemming from the loss of cohesin from the chromosomes during stress, as quantified previously in human cells (Amat et al., 2019). Interestingly, the loss of contacts within TADs was accompanied by an enrichment of contacts between TADs of the *Drosophila* S2 but not in the human A375 cells (Figure 5C and 5D). We suggest that while the depletion of cohesin from human chromosomes leads to the overall decompaction of genomic loci, as observed with microscopy (Luppino et al., 2020), the *Drosophila* genome may depend on alternative mechanisms, potentially facilitated by phase separation through its abundant architectural proteins, to enhance chromatin condensation in the absence of loop extrusion factors (Bing et al., 2024). The recovery condition shows a re-gain of contacts within and between TADs compared to the stress condition in A375 but not in *Drosophila* TADs. These results show that unlike the *Drosophila* S2 cells, the human A375 cells are able to re-form their TADs to levels almost equivalent to those of the unstressed cells within only one hour. In fact, we found that in human cells, control and recovery insulation scores are most correlated (R = 0.9763), showing the effectiveness of recovery, while in S2 cells, the stress and recovery are most correlated (R = 0.9968), showing a lack of reversion to control state (Supplemental Figures S4A and S4B). Despite the substantial re-formation of TADs in A375 recovery cells, boundary strength is not fully restored in either organism. We deduce that the restoration of TADs in the human genome may occur rapidly as cohesin and CTCF return to chromatin, but the establishment of full boundary strength requires processes that are slower to re-establish. The establishment of TADs may proceed even more slowly in *Drosophila*, as neither its TAD count nor boundary strengths were restored following osmotic stress.

### Hyperosmotic stress leads to major loss of small-scale features

DNA regulatory interactions such as enhancer-promoter and promoter-promoter contacts are documented to occur at a finer scale than TADs, with some occurring even at sub-kilobase levels (Eagen et al., 2017; Hsieh et al., 2022; Lajoie et al., 2015). In mammals, such high-resolution contacts can present as isolated dots of high contacts in the contact matrix that often correspond to pairing of convergent CTCF-binding sites (Rowley et al., 2017). Available evidence shows that *Drosophila* domains lack CTCF-cohesin-mediated hotspots (Matthews and White, 2019). Instead, the majority of *Drosophila* domain peaks are said to be derived from the interaction of flanking domains (Bing et al., 2024; Matthews and White, 2019; Rowley and Corces, 2018) and only a subset of dots similar those found in mammals have been described, lacking the typical CTCF signature binding sites (Eagen et al., 2017). However, our Hi-C analysis of the *Drosophila* S2 cells revealed the existence of strong point-to-point interaction hotspots that seem to behave under normal and osmotic stress conditions in a way similar to those found in mammals (Figure 6A).

**Figure 6:**
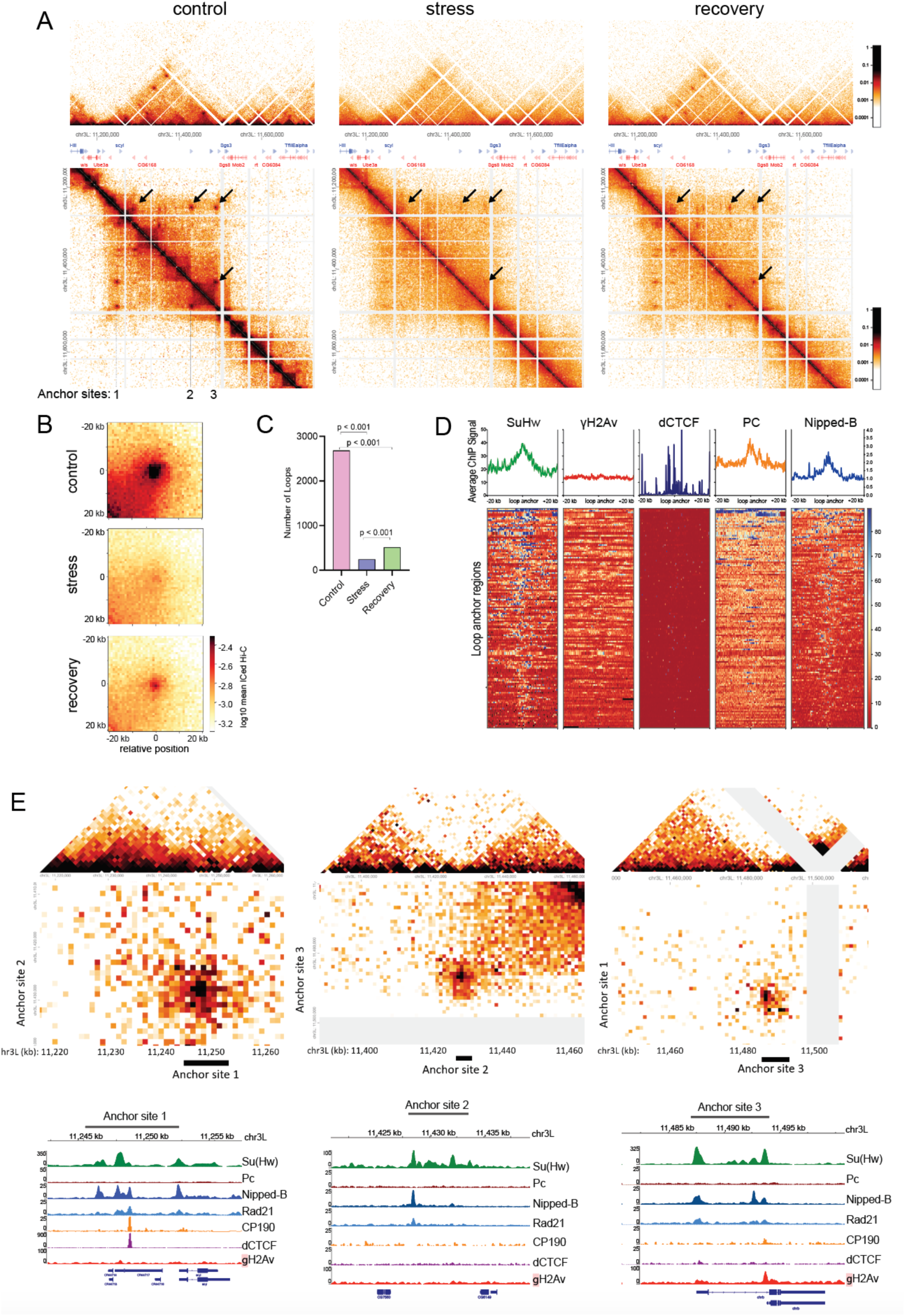
Hyperosmotic stress leads to loss of loops in *Drosophila* cells, but loops occupied by insulator proteins partially recover. A. Comparison of Hi-C heatmaps (2 kb) of *Drosophila* chromosome 3L in control, stress (250mM NaCl) and recovery. Focal points (arrows) correspond to loops that disappear under hyperosmotic stress conditions and reemerge during recovery after osmotic stress. B. Hi-C pileup heatmap averaging the contacts surrounding loops that were detected in control and recovery, but not in stress in *Drosophila* S2 cells. C. Total number of loops in *Drosophila* S2 cells after control, stress and recovery treatments. P values calculated using Fisher’s exact test for each pair of conditions. D. Comparison of *Drosophila* ChIP-seq data of architectural proteins at loop anchors, for loops detected in Control and Recovery, but not stress conditions. Insulator protein Su(Hw), Polycomb (PC) and the cohesin Nipped-B protein reveal enrichment at loop anchor sites. E. 1 kb resolution Hi-C heatmaps at each of the 3 anchors identified in A, showing boundary effects near the diagonal (top triangles) superimposed on looping interaction with another anchor (bottom square). Below each Hi-C heatmap are ChIP-seq profile of insulator proteins, γH2Av, PC and cohesin proteins Nipped-B and Rad-21, at each anchor site

As stated above, the human CTCF and the *Drosophila* insulator proteins display different responses to osmotic stress in that both are released from chromatin but only *Drosophila* insulators form insulator body condensates (Figure 1). Subsequently, osmotic stress in human cells completely abolishes loop structures, which recover upon return to standard media (as seen in Figure 5a, supplemental figure S5 and reported by (Amat et al., 2019). However, considering that loop structures in humans largely rely on CTCF while *Drosophila* loops seemingly do not (Acemel and Lupiáñez, 2023; Eagen et al., 2017; Matthews and White, 2019), we wondered how osmotic stress would affect loop features in *Drosophila*. Loops were called at 5kb resolution and examined using Hi-C heat maps and Hi-C pileup heatmaps (Figure 6A and 6B, respectively). Similar to human cells, most loops in the *Drosophila* S2 cells were lost after osmotic stress. Figure 6A shows loops spanning bases 11,200 kb to 11,500 kb on chromosome chr3L. Loops appear to be formed between all possible combinations of 3 anchor sites. Loops are lost during osmotic stress, but visibly begin to reappear after recovery (Figure 6A). Figure 6B shows a genome-wide Hi-C pileup heatmap of loops that are lost under osmotic stress but re-form after recovery in normal media. We note that loops re-appear more strongly than boundaries created around these loop anchors (see examples in Figure 6A and S5A-E, pileup in 6B). Not all detected loops are regained in this way: 2169, 244, and 511 loops were detected in the control, stress, and recovery conditions respectively (Figure 6C).

For both human and *Drosophila* cells, we interpret the loss of loops during osmotic stress as a direct consequence of the release of cohesin and insulator proteins from chromatin as shown in Figure 1 and as was previously reported (Amankwaa et al., 2022; Amat et al., 2019; Schoborg et al., 2013). In *Drosophila*, however, the formation of loops has been associated with enrichment of Polycomb and Cohesin, but no reported evidence so far indicates the involvement of insulator proteins such as dCTCF (Eagen et al., 2017; Loubiere et al., 2020). We therefore generated enrichment plots for ChIP-seq data at loop anchors to test the enrichment of insulator proteins and other architectural proteins at loop anchor sites (Figure 6D). Insulator protein Su(Hw), Polycomb (PC) and the cohesin subunit Nipped-B revealed enrichment at anchor sites when only the set of loops detected in control conditions, lost in salt stress and recovered after return to normal conditions were used as reference points (Figure 6D). dCTCF, CP190 and γH2Av do not show substantial enrichment in association with the same loop anchors.

Supporting this observation, Figure 6E shows the ChIP-seq profile of insulator proteins, γH2Av, PC, and cohesin proteins Nipped-B and Rad-21 at the three loop anchor sites identified in Figure 6A. Anchor sites appear consistently enriched in Su(Hw) and cohesin, but not in PC, γH2Av, dCTCF or CP190. Figure 6E also shows a magnified display of the dot and anchor site Hi-C heatmaps, revealing that anchor sites span several kb and have boundary activity. Notably, the spread of Su(Hw) and Nipped-B enrichment in the DNA roughly parallels the size of the anchor sites as revealed by Hi-C contact maps.

In supplemental Figure S5, we provide further examples of loop structures and their associated architectural protein enrichment through the *Drosophila* genome. Examples include loops with enrichment profiles in which PC and γH2Av are variable and not always present (Supplemental Figure S5). However all examined instances show significant enrichment in Su(Hw), Nipped-B and Rad-21. Supplemental Figure S5G, for example, shows Nipped-B, Rad-21 and Su(Hw) enrichment that extends over more than 20 kb on chromosome 3L, mediating the formation of a continuous stretch of high frequency long-range loop contacts as well as a small TAD that is produced by these interactions. Unlike interaction domains formed between more distantly spaced loop anchors (as in Figure 6A), where the TAD interactions and boundaries do not recover as quickly as the loop, this smaller interaction domain that is completely coated by insulators shows rapid recovery. Considering the currently available data, we conclude that these structures form in the absence of any significant contribution from PC or other architectural proteins such as dCTCF or CP190. These observations suggest Su(Hw) and cohesin play a critical role in loop formation in the *Drosophila* genome. However, not all insulator-enriched sites form loops. Notably, Supplemental Figure S5E illustrates a case where a TAD boundary shows strong enrichment in Su(Hw), cohesin and γH2Av yet does not form a loop and does not recover after return to normal media. This dissociation between TAD boundary activity and loop formation demonstrates that these two structural features, while often co-occurring, are mechanistically distinct in *Drosophila*.

### γH2Av and Su(Hw) recapitulate *Drosophila* compartment identity

In mammals, compartments are typically seen in the range of 100kb. However, similar to an earlier report (Rowley et al., 2017), our analysis revealed compartments at a resolution as high as 2kb in the *Drosophila* genome (Figure 7A). Just like compartments in humans, we found that the A compartments in the *Drosophila* genome engage in a long-range A-to-A interactions (Figure 7A). Surprisingly, this high-resolution compartmentalization in *Drosophila* appears to deviate from the traditional checker-board patterns originally found in the human genome. First, unlike the A375 cells in which the A and B compartments alternate throughout the genome (Figure 3), we observed that in *Drosophila*, most B compartments do not engage in strong long-range B-to-B interactions (Figure 7A). This lack of strong B-to-B interactions in the long-range is also confirmed using saddle plot analysis, where B-B interactions are much less extensive than A-A interactions (Figure 4C).

**Figure 7:**
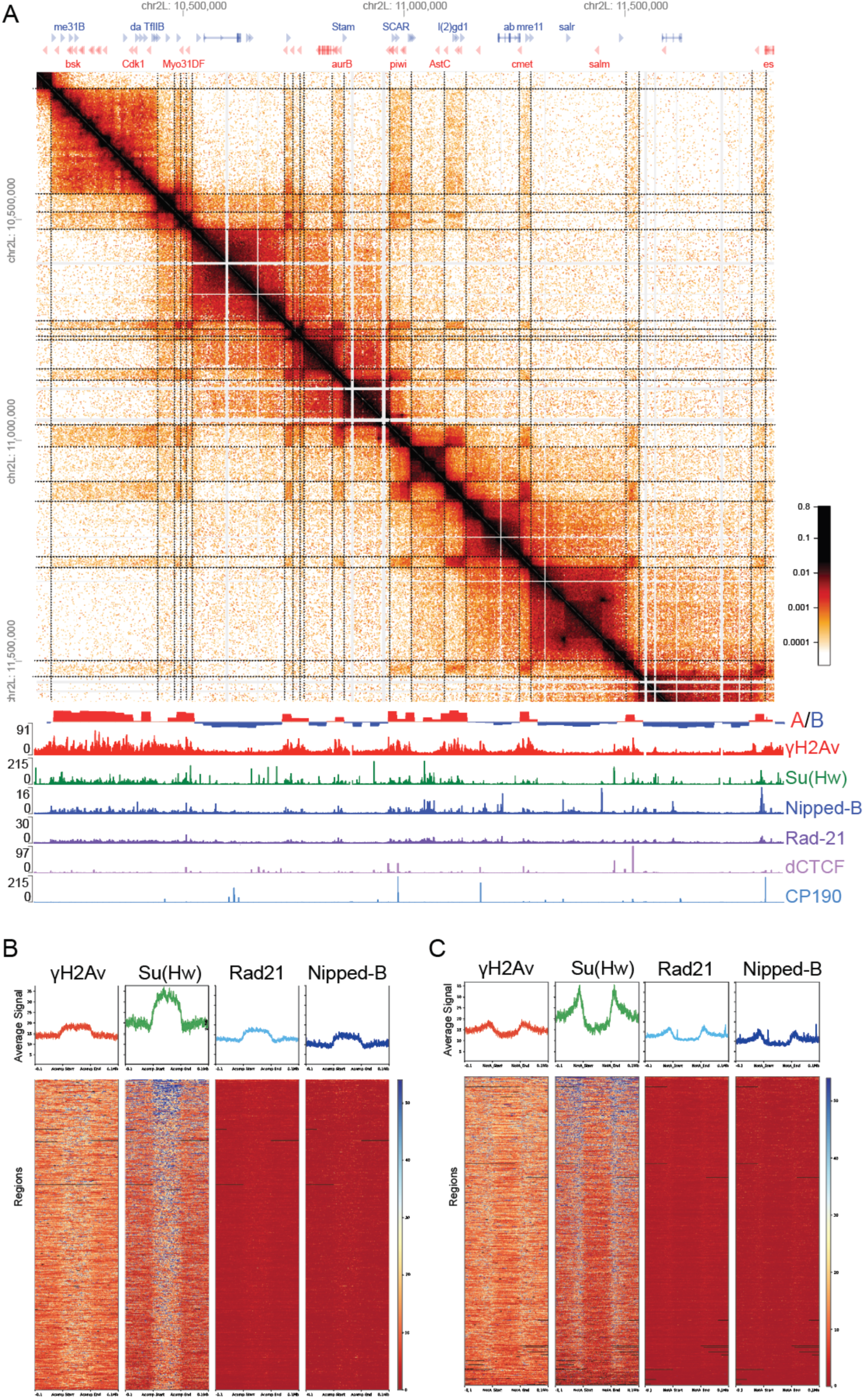
A compartments in *Drosophila* align with γH2Av, cohesin, and Su(Hw) domains. **A**. “A” compartments in normal media (control) engage in long-range interactions with other “A” compartments. “B” compartments do not engage in long-range interactions. Dotted lines indicate compartment boundaries. Below the heatmap is seen the ChIP-seq profile distribution of insulator proteins showing “A” compartments are enriched in γH2Av and Su(Hw) and some cohesin subunits, but poorly enriched in other insulators. This association occurs across the genome. **B**, **C**. ChIP-seq profile distribution of insulator proteins showing “A” compartments are enriched in insulators (B) while “B” compartments are poorly enriched in the insulators (C).

We sought to explore the architectural proteins associated with this unique compartment organization and whether these proteins contribute to the differences in domain restructuring following osmotic stress and recovery compared to the response by the human genome. We therefore aligned publicly available ChIP-seq data of insulator associated proteins such as Su(Hw), CP190, dCTCF, γH2Av, and the cohesin proteins, Rad21, Wapl, and Nipped-B to our control *Drosophila* Hi-C data. See Methods for sources of all datasets used.

We found that within this set of proteins, the A compartments (called at 20 kb resolution) in the *Drosophila* genome were enriched for γH2Av and Su(Hw) (Figure 7B). Additionally, we observed a moderate enrichment of the cohesin subunits Rad21, and Nipped-B at the A-compartment sites (Figure 7 B and C). This is interesting against the backdrop of the genome-wide close association between Su(Hw) and γH2Av (Amankwaa et al., 2022; Simmons et al., 2022). On the contrary, γH2Av and Su(Hw) were depleted in the non-A (B compartment) regions (Figure 7C). These results suggest that Su(Hw), γH2Av and cohesin may be synergistically involved in *Drosophila* genome compartmentalization. Because *Drosophila* architectural proteins such as γH2Av and the cohesin subunits have the capacity to form condensates (Amankwaa et al., 2022), we deduce that their possible involvement in compartmentalization may occur via liquid-liquid phase separation.

We note that while our Hi-C was performed in S2 cells, some ChIP-seq datasets used in this analysis were generated in other Drosophila cell lines including Kc167 and BG3 cells. Importantly, the consistency of γH2Av and Su(Hw) enrichment patterns at A compartment regions across these independently generated datasets from different cell lines argues that this association reflects a general feature of Drosophila genome organization rather than a cell-type specific property

## Discussion

Owing to advances in genomic techniques, significant progress has been made in understanding the 3D network of chromosome contacts and how it impacts gene expression. It is now widely accepted that the active loop extrusion process and compartmentalization are the primary drivers of the 3D genome. However, there remain several unresolved questions, such as why certain mammalian TADs do not conform to the loop model. Moreover, despite harboring multiple architectural proteins, the *Drosophila* genome appears to be structured through principles that deviate from the loop extrusion model. This has led to the view that the *Drosophila* contact domains may be organized mainly through phase separation. Strikingly, in response to hyperosmotic stress, *Drosophila* IBPs and cohesin demix to form phase separated condensates (Amankwaa et al., 2022; Schoborg et al., 2013). In this work, we sought to harness the unique osmo-sensitive nature of architectural proteins to gain insights into the genome organization principles in *Drosophila* and to understand how these principles differ from those in the human genome.

Our analysis shows that both the *Drosophila* and human genomes undergo significant condensation when exposed to hyperosmotic stress (Figure 2). Osmotic stress-induced chromatin condensation is a well-documented phenomenon, and it is suggested to be a protective mechanism to maintain structural genome integrity (Irianto et al., 2013; Majumder and Jain, 2020; Olins et al., 2020). The converse appears to also be true as isolated chromosome decondensation occurs in low ionic strength conditions (Poirier et al., 2002; Sanders et al., 2022). Thus, the comparable adjustments in chromatin size in response to osmotic stress suggest a conserved mechanism aimed at safeguarding genome integrity. Whereas the *Drosophila* IBP complexes formed condensates after osmotic stress, the human CTCF did not (Figure 2). This is surprising given that hCTCF is predicted to possess LLPS features similar to the *Drosophila* IBPs (Amankwaa et al., 2022), and has been shown to exhibit moderate sensitivity to 1,6-hexanediol, a phase separation-sensitive chemical (Shi et al., 2021). Our extrapolation is that insulator body formation is a manifestation of an intrinsic phase separation behavior of *Drosophila* insulator proteins, a property amplified during osmotic stress.

Despite the differences in condensate behavior between Drosophila and Human insulator proteins, we observed an overall loss of short-range interactions but a gain of long-range interactions in both species after osmotic stress. The loss of local contacts can be attributed to the loss of cohesin, a factor crucial for bringing together nearby genomic segments, along with other proteins that aid in forming loops and facilitating local phase-separating interactions (Schwarzer et al., 2017). Meanwhile, the global condensation of the chromosomes observed using microscopy creates the generic consequence of chromatin regions separated by longer distances being brought together on average. This idea is echoed in the observation that osmotic stress induces the disruption of domain boundaries (Figure 4, Figure 5 and (Amat et al., 2019). The loss of domain boundaries likely results in random mingling of normally spatially segregated local *loci* and a general loss of homotypic interactions for heterotypic contacts as shown in Figures 4 and 5. Interestingly, the removal of boundary proteins such as CTCF or factors such as cohesin alone in other systems is not enough to disrupt A/B compartment interactions (Nuebler et al., 2018; Rao et al., 2017), emphasizing the dichotomy between loops and TADs in *Drosophila* and raising the question of what mechanisms maintain compartment boundaries in mammals in the absence of cohesin (Mirny et al., 2019; Narendra et al., 2016).

The osmotic stress response further reveals a mechanistic decoupling between loops, TAD boundaries and compartments in *Drosophila*. In *Drosophila*, disrupting architectural protein occupancy simultaneously affects all three levels of genome organization (loops, TAD boundaries and compartments) in ways not observed upon removal of individual factors in mammalian systems, underscoring the integrated nature of insulator-dependent genome organization in flies. The observation that certain TAD boundaries enriched in insulator proteins fail to form loops and fail to recover after stress (Supplemental Figure S5E) further supports this decoupling, suggesting that insulator occupancy alone is insufficient for loop formation and that additional factors or configurations are required.

A number of reasons could be given for the distinct osmotic stress recovery patterns between *Drosophila* and human genomes. Studies in mammalian cell cultures indicated that loop extrusion exhibits greater dynamism in comparison to compartmentalization, which is thought to occur through passive mechanisms like phase separation (Oudelaar and Higgs, 2021; Zhang et al., 2019). For instance, relative timing of TAD and compartment re-establishment after mitosis, showed that TADs and cohesin-mediated CTCF-CTCF loops emerge faster than long-range compartmentalization (Abramo et al., 2019). We therefore expect differences in recovery patterns if there exists a substantial difference in how loop extrusion and compartmentalization contribute to shaping the *Drosophila* and mammalian genomes. In human A375 cells, the chromatin size in recovery cells is restored to a level comparable to the pre-osmostress state (Figure 1B), whereas in S2 cells recovery remains significantly below the original control level (Figure 1D).

The contribution of loop extrusion to loop formation in *Drosophila* is considered negligible, given the lack of evidence implicating insulator proteins in the process of loop formation in the *Drosophila* genome (Eagen et al., 2017; Li et al., 2023). In *Drosophila*, loops do not result from the distinct interaction between cohesin and CTCF at loop anchors. Instead, *Drosophila* loop anchor sites have been described as enriched in Polycomb (PC) and cohesin but not in insulator proteins (Eagen et al., 2017; Loubiere et al., 2020).

One striking result from our work is that *Drosophila* loops dissolve during osmotic stress and reorganize after recovery in normal media, a behavior that mirrors loops in the human genome. This response also parallels that of insulator proteins in both human and *Drosophila*, as they exit chromatin under osmotic stress and bind back to chromatin after recovery (Amat et al., 2019; Schoborg et al., 2013). ChIP-seq analysis of insulator proteins revealed that the insulator protein Su(Hw) is enriched at loop anchor sites and that these sites show boundary activity (Figure 6 and supplemental Figure S5). It is intriguing that PC does not exhibit the same response as insulator proteins to osmotic stress (Schoborg et al., 2013), and that not all loop anchors examined in this work contain PC (Figure 6 and Supplemental Figure S5). Instead, anchor sites are enriched in Su(Hw) plus cohesin, which is remarkably similar to how CTCF and cohesin are found at loop anchors in humans. These observations raise the possibility that Su(Hw) and cohesin cooperate to mediate loop formation in *Drosophila* through a mechanism that, while distinct from canonical CTCF-cohesin loop extrusion, similarly depends on the co-occupancy of a sequence-specific DNA binding protein and cohesin at loop anchors. Whether this represents a loop extrusion-like mechanism or is better explained by the boundary pairing model recently proposed for *Drosophila* domain organization (Bing et al., 2024) remains an important open question.

Given the similarity in the response of loops after exposure to osmotic stress, the unique way in which the human and *Drosophila* genomes undergo compartment restructuring is striking. Interestingly, saddle plots reveal that human A-A interactions do not fully re-establish after recovery from stress (Figure 4). Given that the *Drosophila* genome largely depends on A-A interactions, this suggests that the genomic organization in *Drosophila* and humans might not be as different as it looks at first glance. The more efficient re-organization of compartments observed in humans could be explained because they are more heavily driven by B-B interactions (Falk et al., 2019). Specifically, while both A-A and B-B interactions are lost in the human genome, osmotic stress appears to induce a preference for more B-B interactions in the *Drosophila* cells (Figure 4). It is also possible that the increase in B interactions observed in *Drosophila* involves regions with weak A and weak B interactions that were not strongly compartmentalized to begin with. This finding is intriguing because, in contrast to the human genome, specific compartments within *Drosophila* control cells did not alternate to create the typical A and B compartment checkerboard patterns (Figure 3 and Figure 7).

The mechanism responsible for this unusual compartment arrangement is not clear. However, it is obvious that over long distances, the *Drosophila* B compartments seldom participate in B-B interactions (Figure 2.3). A similar observation has recently been made by others (Harris et al., 2023; Rowley et al., 2019). In particular, Rowley *et al* posited that there is a drastic difference in stronger A-A interactions than B-B interactions in the *Drosophila* genome (Rowley et al., 2019). In the same work, it was demonstrated that in the *Drosophila* genome, RNA polymerase II within gene bodies is critical for strong A-A interactions (Rowley et al., 2019). These may contribute to the apparent lack of long-range B-B contacts.

Out of the chromatin architecture proteins we analyzed, the *Drosophila* phosphorylated H2A variant (γH2Av) and Su(Hw) most closely reflected eigenvector changes, showing enrichment in the A compartments and depletion in the B compartments. It has been shown that reduced deposition of H2Av at gene promoters, through a knockdown of the histone chaperone Domino, disorganizes the *Drosophila* 3D genome (Ibarra-Morales et al., 2021) and that eigenvector transitions coincided closely with H2Av at 10 kb resolution (Rowley et al., 2017). These reports do not clarify whether the impact on the 3D genome can be attributed to the phosphorylated or unphosphorylated forms of H2Av. However, we showed earlier that phosphatase inhibition stabilizes the interactions between γH2Av and gypsy insulator proteins and that γH2Av and Su(Hw) are enriched at TAD boundaries (Simmons et al., 2022). Interestingly, the phosphorylated (γH2Av) and not the unphosphorylated H2Av undergoes phase separation together with the canonical *Drosophila* IBPs (Amankwaa et al., 2022; Simmons et al., 2022).

Based on these findings, we deduce that the role of γH2Av goes beyond being a gene expression and DNA damage response protein like its human homologs. Rather, it appears to be a *Drosophila* specific chromatin state and compartment dictating modification.

Notwithstanding these Drosophila-specific features, the incomplete recovery of genome organization we observe in both species raises a broader question about the biological consequences of transient chromatin reorganization. Although human genome structure recovered far more completely than *Drosophila* after osmotic stress, even transient 3D genome organization changes may have a lasting biological impact. Osmotic forces arising from cell compression have been shown to alter chromatin organization and stem cell fate (McCreery et al., 2025), suggesting that stress-induced contact changes are physiologically relevant beyond the experimental context. The new contacts formed during stress and the compartment and TAD changes that fail to recover in our system, could therefore underlie longer-term phenotypic effects of osmotic stress on genome regulation.

Together, our results show that both the human and *Drosophila* genomes display similar responses to hyperosmotic stress, characterized by chromatin shrinkage, exit of architectural proteins from the chromatin, loss of domain boundaries, loss of loops and overall loss of short-range but a gain of long-range contacts. However, both organisms display certain distinct genome restructuring responses following the osmotic stress. Notably, in contrast to *Drosophila*, the human genome demonstrates a more robust or rapid restoration of lost genome structures induced by osmotic stress. We propose that the slower recovery of *Drosophila* genome organization reflects a fundamental difference in the mechanisms maintaining compartment structure in the two species. In humans, TAD reformation is driven largely by loop extrusion, a fast and mechanistically simple process that requires only the re-loading of cohesin onto CTCF-bound chromatin. Compartmentalization, which depends on slower phase-separation-like mechanisms in both species, recovers more slowly in both, as evidenced by the incomplete restoration of A-A interactions in human recovery cells (Figure 4D). In *Drosophila*, where compartment organization depends predominantly on γH2Av and Su(Hw) rather than on loop extrusion, the absence of a fast loop-extrusion-driven recovery pathway means that both TADs and compartments must await the slower re-establishment of concentration-dependent interactions and possibly LLPS, consistent with the near-complete lack of structural recovery we observe.

## Materials and Methods

### S2 Cell Culture

Cells were maintained in Schneider’s *Drosophila* Media (1X) (Gibco) supplemented with 10% fetal bovine serum to form complete Schneider’s *Drosophila* Media at 25°C.

### Human Cell Culture

A375 cells were cultured in Dulbecco’s Modified Eagle Medium (DMEM, Corning), supplemented with 10% fetal bovine serum, 1% Penicillin-Streptomycin, and 1% L-Glutamine, at 37°C in a 5% CO_2_ atmosphere.

### Hyperosmotic Stress Treatment and Immunostaining

S2 cells were sub-cultured for 3 days at 25°C. Prior to the immunostaining process, the cells were seeded onto a poly-l-lysine-treated coverslips and allowed to adhere for 30 minutes in a covered 35-mm cell culture dish. Osmotic stress was induced in the cells by quickly replacing the conditioned media with fresh indicated concentration of NaCl (from a 5M stock) prepared with complete Schneider’s *Drosophila* media while the control cells were kept in the conditioned media. To induce stress, the cells were kept in the specified concentration of NaCl for 1 hour, following by immunostaining as previously described (Rogers and Rogers, 2008).

In summary, the S2 cells were fixed with 4% PFA for 10 minutes at room temperature (RT), followed by three PBS washes. The fixed cells were permeabilized with 0.2% Triton X-100 for 5 minutes followed by blocking with 3% nonfat milk for 10 minutes at RT. Primary antibodies were prepared by diluting an aliquoted stock in 3% nonfat milk. The cells were incubated in the primary antibody for 1 hr at RT in a humidified chamber followed by a 3X wash with PBS/0.1% Triton X-100 for 5 min each. This was followed by a 1hour incubation in a secondary antibody prepared from a 3% nonfat milk. Subsequently, the coverslips were washed as described for the primary antibody staining. To counterstain DNA, coverslips were treated with 0.5 μg/ml DAPI, rinsed twice with H_2_O, and mounted in Vectashield.

On the other hand, prior to the experiment, we sub-cultured the A375 cells on flame sterilized coverslips in a 6-well dish for 3 days prior to experiment. Osmotic stress was induced by replacing the media with fresh one containing required concentrations of NaCl prepared from a 5M stock. The stressing was done for 1 hour at 5% CO_2_ and 37°C. At the same time, control cells were kept in fresh DMEM. We then fixed the cells with 4% PFA for 15 minutes at room temperature (RT), followed by three PBS washes, permeabilization with 0.25% Triton-X100 for 5 minutes. The permeabilization step was followed by primary antibody staining overnight in a blocking buffer at 4°C. We did secondary antibody staining in the dark with blocking buffer diluted antibody solution for 1 hour. This was followed by 2X washes with cold DPBS. To stain DNA, coverslips were treated with 0.5 μg/ml DAPI, rinsed twice with H2O, and mounted in Vectashield.

### Antibodies

anti -dCTCF from guinea pig [gift from Elissa Lei]. And anti-human CTCF antibody, purchased from Abcam (ab128873) were used for immunostaining, used at a final dilution of 1:100 and 1:200 respectively.

### Fluorescence and confocal microcopy

All microscopy for immunostaining was performed on a wide-field epi-fluorescent microscope (DM6000 B; Leica Microsystems) equipped with a 100×/1.35 NA oil immersion objective and a charge-coupled device camera (ORCA-ER; Hamamatsu Photonics). Simple PCI (v6.6; Hamamatsu Photonics) was used for image acquisition. FIJI, an open-source image-processing package based on ImageJ2 was used for image analysis (Schindelin et al., 2012). All contrast adjustments are linear. Images were further processed in Adobe Photoshop CS5 Extended version 12.0 ×64 and then assembled with Adobe Illustrator version 28.2, Python version 3.7 and GraphPad Prism version 9.0.0 (224) (GraphPad Software) were used to perform the statistical analyses. Only the most typical cases of cytological localizations are shown on the figures in the manuscript in the “Results” section. However, the conclusions are drawn on the basis of analysis of large numbers of cells collected in triplicates.

### Genome-wide Chromosome Conformation Capture (Hi-C)

15 million S2 cells were used for each of the two replicates for each condition. Prior to the Hi-C assay, the S2 cells were pretreated with just the complete Schneider’s media and 250mM NaCl prepared in complete Schneider’s media as control and stress respectively for 1 hour. The recovery cells underwent the same stress treatment for 1 hour, followed by a 1 hour recovery in the complete Schneider’s media. After these treatments, the cells were strained passing them through a 40µm strainer to remove clumps. We then fixed the cells with 2% formaldehyde at RT for 10 minutes. The crosslinking was quenched with 125mM glycine. Following fixation and quenching, nuclei were isolated using 5 mL cold lysis buffer (50 mM Tris-HCl pH 7.5, 150 mM NaCl, 5 mM EDTA, 0.5% NP-40, 1% Triton, 1x Roche Complete Protease Inhibitors). The isolated nuclei were then snap frozen in liquid nitrogen and stored at −80°C for later Hi-C analysis.

The human A375 malignant melanoma cells were purchased from ATCC (ATCC® CRL-1619™). The cells were grown in DMEM enriched with 10% FBS, 1% Pen-strep, and 1% L-glutamine, and were sub-cultured to at a density of 80%. For salt stress experiments, A375 cells were introduced into media supplemented with 250mM NaCl for one hour, for stress samples, or subjected to 250mM NaCl for one hour followed by reintroduced to conditioned DMEM for an additional hour, for recovery samples.

For both *Drosophila* S2 and human A375 cells, Hi-C was performed using the Arima Hi-C (Arima Genomics) kit, following the manufacturer’s protocol A160134 v01 for library amplification using the NEBNext Ultra II kit (NEB;E7645S). Sequencing was performed by Genewiz on either an Illumina NovaSeq or HiSeq platform with 50 or 150 bp paired-end reads. The sequenced reads were aligned to the reference human and *Drosophila* genomes hg19 and dm6 respectively, binned, and iteratively corrected following the hic-pro mapping pipeline accessible at https://github.com/nservant/HiC-Pro.

### Distance Decay Scaling Plots

The global distance decay plots at the territorial level were calculated using calculated using the cooltools suite https://cooltools.readthedocs.io/en/latest/notebooks/contacts_vs_distance.html The average contact frequency at each distance genome-wide was calculated and plotted vs. genomic distance on a log scale.

### Compartment Analysis

Principal component analysis using matrix2compartment.pl script in the cworld-dekker pipeline available on GitHub (https://github.com/dekkerlab/cworld-dekker) was used for compartment analysis. The human and *Drosophila* genome compartment identity was determined for 250Kb and 20kb binned matrices respectively, using the first eigenvector values (PC1). Positive and negative PC1 values were respectively denoted A and B compartments, where positive values were assigned by the script based on correlation with overall gene density. Compartment switches were classified as those bins that changed sign (negative to positive for B to A or positive to negative for A to B) and also shifted value by at least 10% of the total PC1 range. Compartment switches were classified as “fully recovered” if their value in the Recovery condition was within 2% of the PC1 value in Control or “not recovered” if the PC1 value was still within 2% of the Stress condition. Everything in between was classified “partially recovered”.

### Saddle Plots

Saddle plots were used to analyze homotypic and heterotypic A and B interactions. Using the matrix2compartment.pl script in cworld-dekker, we calculated interaction Zscores, normalizing for interaction decay over genomic distance. The Zscore matrices were reordered from more negative to positive compartment eigenvector values signifying strongest B to strongest A compartments. Finally, matrices were smoothed at 500 kb for A375 and 40 kb for S2 cells. Compartment strength was calculated using the saddle plot functions in coolTools (Open2C et al., 2024) (https://cooltools.readthedocs.io/en/latest/notebooks/compartments_and_saddles.html). The bins with the strongest 20% of positive (for A) or negative (for B) PC1 values were used to calculate each interaction category in the equation Compartment strength = (AA+BB)/AB.

### *Drosophila* ChIP-seq data

Previously published ChIP-seq datasets for *Drosophila* insulators and other chromatin binding proteins were used for comparison to Hi-C data features. Datasets used included: Polycomb (S2 cells, SRR26350370), (Brown et al., 2024), dCTCF (Kc cells, SRR317182) (Wood et al., 2011), γH2Av (Kc167 cells, SRR2546860 (Li et al., 2016), Su(Hw) (S2 cells, SRR121539) (Chen et al., 2012), Wapl (BG3 cells, SRR6497804) (Misulovin et al., 2018), Rad21 (BG3 cells, SRR7689010) (Pherson et al., 2019), and Nipped-B (BG3 cells, SRR7688997) (Pherson et al., 2019). Fastq files were trimmed to remove adapters and low quality sequences using trimmomatic (Bolger et al., 2014), and then mapped to the dm6 genome using bowtie2 (Langmead and Salzberg, 2012). The bamCoverage function of deeptools 3.1.1 (Ramirez et al., 2014) was then used to convert mapped .bam files into the bigWig format for visualization.

### Loop calling and Hi-C pileup heatmaps

Isolated dots of highly enriched contacts (“loops”) were detected in the *Drosophila* 5 kb binned Hi-C data using the Mustache tool (Roayaei Ardakany et al., 2020); https://github.com/ay-lab/mustache) with the following parameters: pThreshold = 0.05, sigmaZero = 0.6, and sparsityThreshold = 1.0. Only loops with anchors less than 300 kb away from each other were considered in downstream analyses. To create average Hi-C contact maps around these detected loops, we used the coolTools pileup function; https://cooltools.readthedocs.io/en/latest/notebooks/pileup_CTCF.html) using the 1 kb resolution Hi-C contact maps.

### Enrichment plots for ChIP-seq data at loop anchors and compartments

Enrichment heatmaps and average profile plots were constructed using the computeMatrix, plotProfile and plotHeatmap tools that are part of the deepTools suite on Galaxy (Ramirez et al., 2014). For loop anchor profiles, the set of loops detected in Control conditions, lost in salt stress, and recovered after return to control conditions were used as reference points, with 20 kb upstream and downstream sequence included in each profile. Only loops with anchors less than 300 kb away from each other were considered. For comparison to A and B compartment regions, we classified A compartment regions as regions with strong positive first eigenvector values (>0.025) from PCA on 20 kb contact maps. All other regions were considered B compartment regions. This threshold was necessary because the A compartment regions show much stronger distal interactions than intervening B regions, so the first eigenvector is skewed toward positive values. Each A and B compartment region was scaled to a relative size of 100 kb and then 100 kb upstream and downstream were included as flanking sequences. For both types of enrichment heatmaps, the ChIP-seq data bigwig files were binned into 50 bp bins and the mean value was used for region sorting and average profile plots.

## Data Availability

All raw and processed Hi-C data is available at GEO accession number GSE275816. Publicly available datasets used in analyses are cited in the Methods section.

## Author Contributions

Conceptualization: M.L. and R.P.M. Data Curation: R.P.M. Formal Analysis: B.A.; R.P.M. M.L. and T. X. Funding Acquisition: R.P.M. and M.L. Investigation: B.A., C.P., E.S. and J.S. Methodology: B.A., M.L. and R.P.M. Project Administration: M.L. and R.P.M. Resources: M.L. and R.P.M. Software: B.A., R.P.M and T.X. Supervision: M.L. and R.P.M. Visualization: B.A. M.L. R.P.M. and T.X. Writing - Original Draft: B.A. Writing - Review & Editing: M.L. and R.P.M.

## Acknowledgments

This work was supported partially by US Public Health Service Award from the National Institutes of Health (MH108956) to M.L. and NIGMS award R35GM133557 to RPM with additional support from the College of Arts and Sciences and the Department of Biochemistry and Cellular and Molecular Biology at The University of Tennessee, Knoxville.

## Supplemental Material

**Supplemental table 1:** Hi-C summary statistics.

| Sample | Total Raw Reads | %Both Sides Mapped | %Dangling Ends | Valid Pairs | Unique Valid Pairs | %Cis |
| --- | --- | --- | --- | --- | --- | --- |
| Drosophila-Control-R1 | 272,652,876 | 41.59 | 1.01 | 104,818,095 | 69,612,995 | 96.06 |
| Drosophila-Control-R2 | 378,843,202 | 43.62 | 6.21 | 132,587,684 | 64,851,092 | 96.62 |
| Drosophila-250mMNaCl-R1 | 350,259,769 | 39.41 | 5.21 | 124,996,611 | 77,530,062 | 77.84 |
| Drosophila-250mMNaCl-R2 | 393,876,288 | 39.01 | 1.34 | 144,089,579 | 92,369,832 | 85.2 |
| Drosophila-Recovery-R1 | 430,841,711 | 39.4 | 5.14 | 154,455,818 | 98,016,264 | 78.83 |
| Drosophila-Recovery-R2 | 342,715,233 | 39.41 | 2.37 | 123,354,943 | 76,058,870 | 82.3 |
| OsmoStress-A375-Control-R1 | 164,104,030 | 69.4 | 0.72 | 112,216,528 | 78,245,269 | 76.05 |
| OsmoStress-A375-Control-R2 | 328,064,727 | 61.27 | 6.03 | 184,728,292 | 125,011,691 | 78.72 |
| OsmoStress-A375-250mMNaCl-R1 | 183,751,848 | 68.68 | 1.05 | 123,481,557 | 90,462,782 | 68.07 |
| OsmoStress-A375-250mMNaCl-R2 | 346,454,641 | 60.08 | 6.12 | 190,661,389 | 128,760,880 | 77.35 |
| OsmoStress-A375-Recovery-R1 | 116,635,178 | 69.47 | 1.01 | 79,334,864 | 49,526,110 | 73.71 |
| OsmoStress-A375-Recovery-R2 | 459,891,066 | 62.18 | 4.79 | 266,901,842 | 174,484,556 | 77.33 |

**Supplemental Figure S1:**
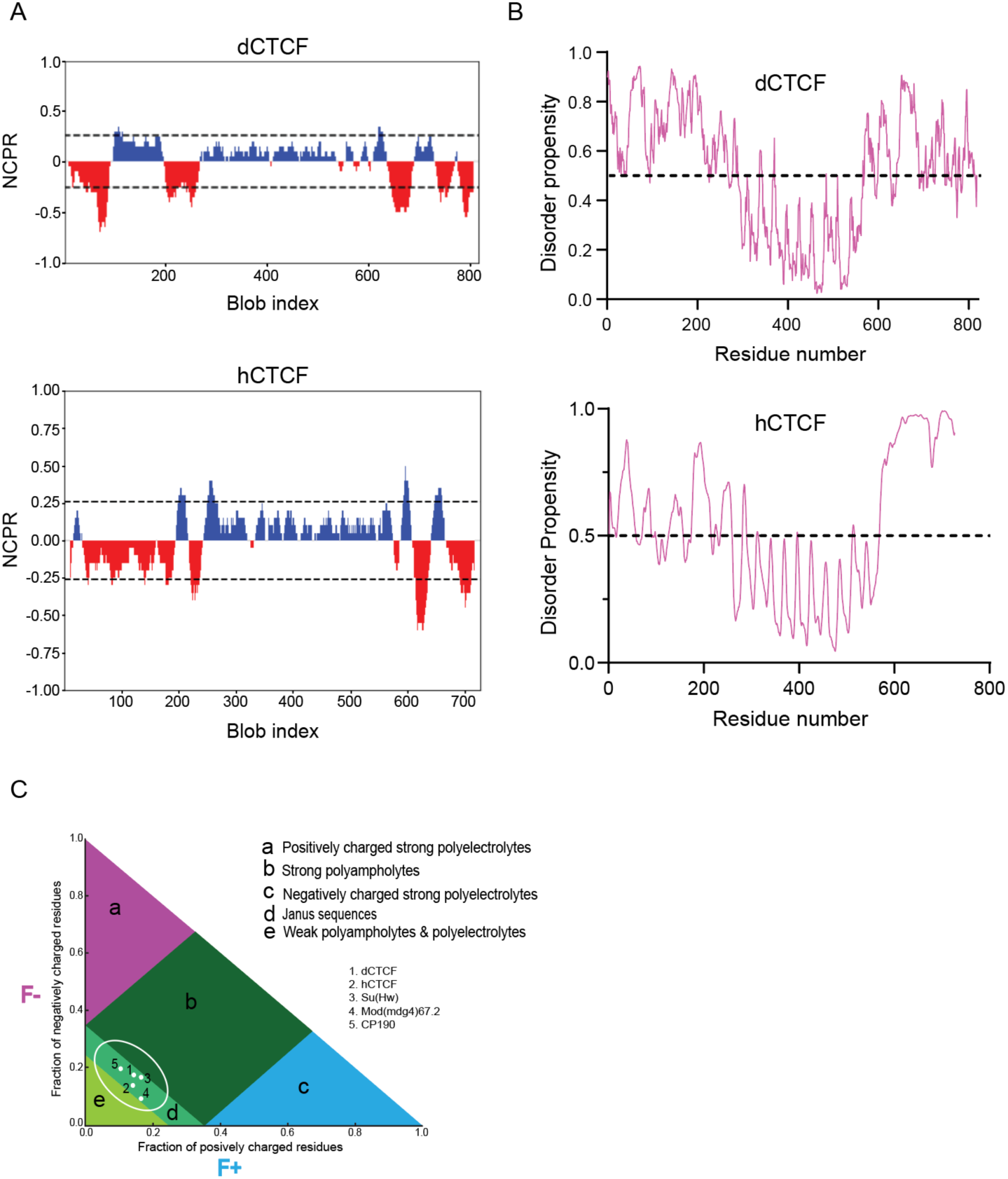
*Drosophila* IBPs and hCTCF share Similar Predicted LLPS Properties. **A**. Analysis of charged versus uncharged amino acid distributions for dCTCF (top panel) and hCTCF (bottom panel). Blue and red peaks indicate positive and negative charged residues respectively while gaps denote non-charged residues. **B**. Disorder tendency comparison between dCTCF and hCTCF. Pink lines above a 0.5 disorder propensity score (Y-axis) denote disordered regions while those below indicate structures domains. **C**. Das-Pappu’s phase diagram showing likely insulator protein disordered conformations. F(−), fraction of negatively charged residues; F (+), fraction of positively charged residues. Protein sequences in regions “a” and “c” depict strong polyelectrolyte features with FCR > 0.35 and net charge per residue > 0.3. Such proteins mostly exhibit coil-like conformations. Region “b” corresponds to strong polyampholytes that form distinctly non-globular conformations, such as coil-like, hairpin-like, or hybrids. Region “e” relates to either weak polyampholytes or weak polyelectrolytes that form globule or tadpole-like.

**Supplemental Figure S2:**
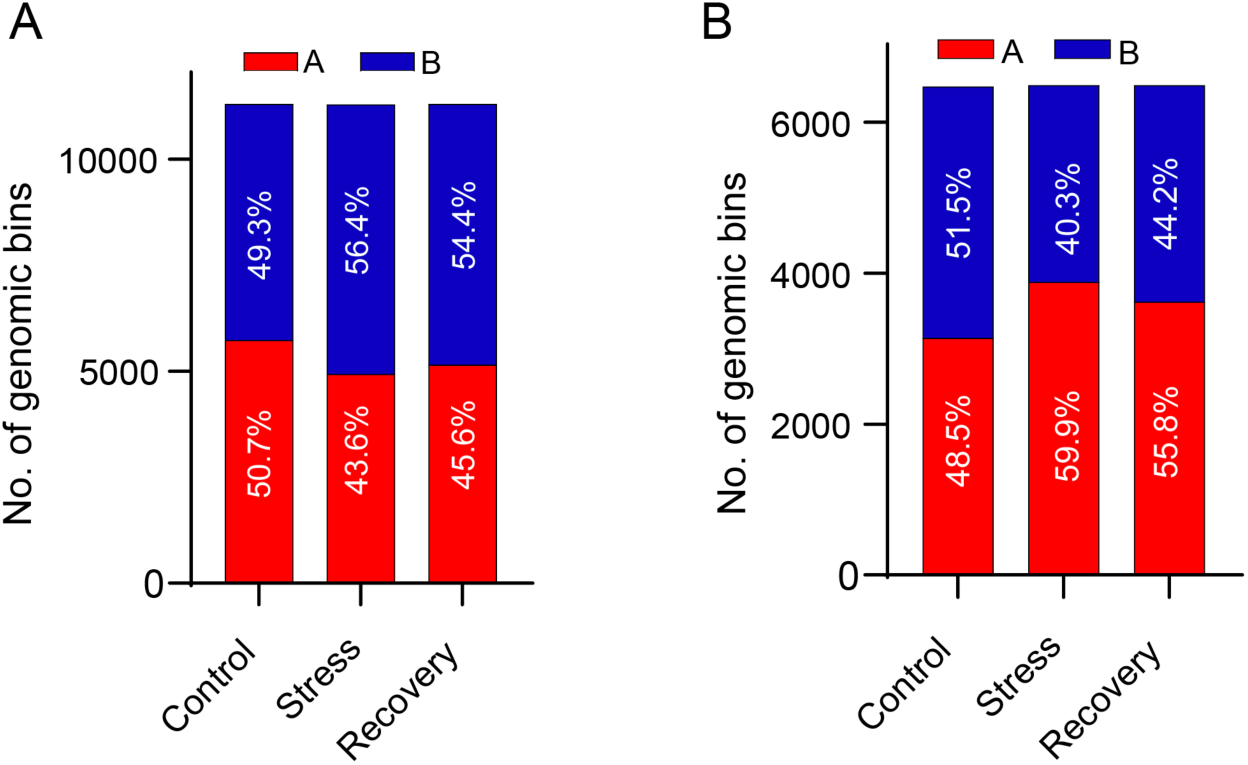
The human A375 and *Drosophila* S2 cell genomes display different proportions of A/B compartment identities after osmotic stress and recovery. **A**. Proportions of A (red) and B (blue) compartments defined by first principal component values in control, osmotic stress, and recovery of the human A375 cells. **B**. Proportions of A (red) and B (blue) compartments defined by first principal component values in control, osmotic stress, and recovery of *Drosophila* S2 cells.

**Supplemental Figure S3:**
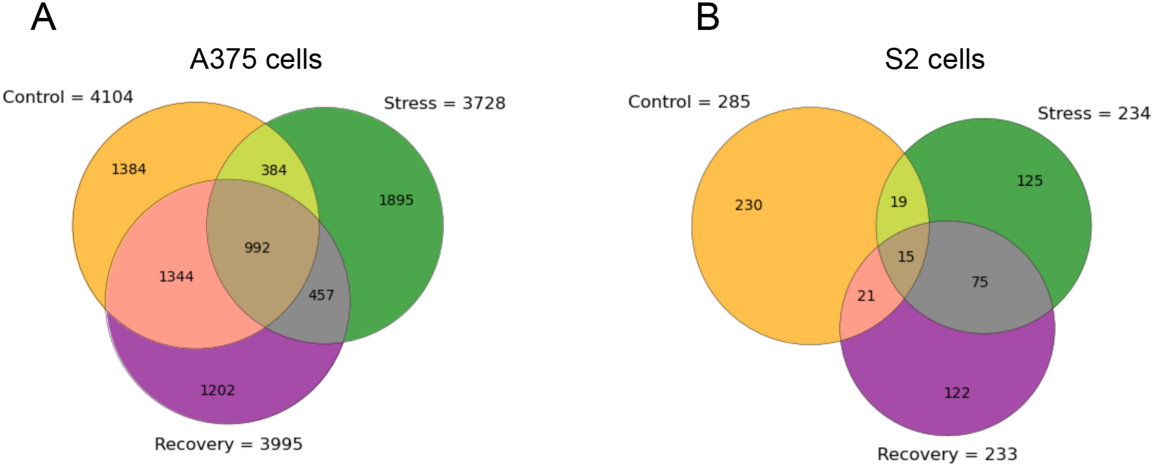
Human A375 and *Drosophila* S2 Cells show differences in TAD boundary changes after osmotic stress and recovery. **A**. TAD boundary number stratifications in human A375 controls, stress, and recovery. **B**. TAD boundary number stratifications in *Drosophila* S2 cell controls, stress, and recovery.

**Supplemental Figure S4:**
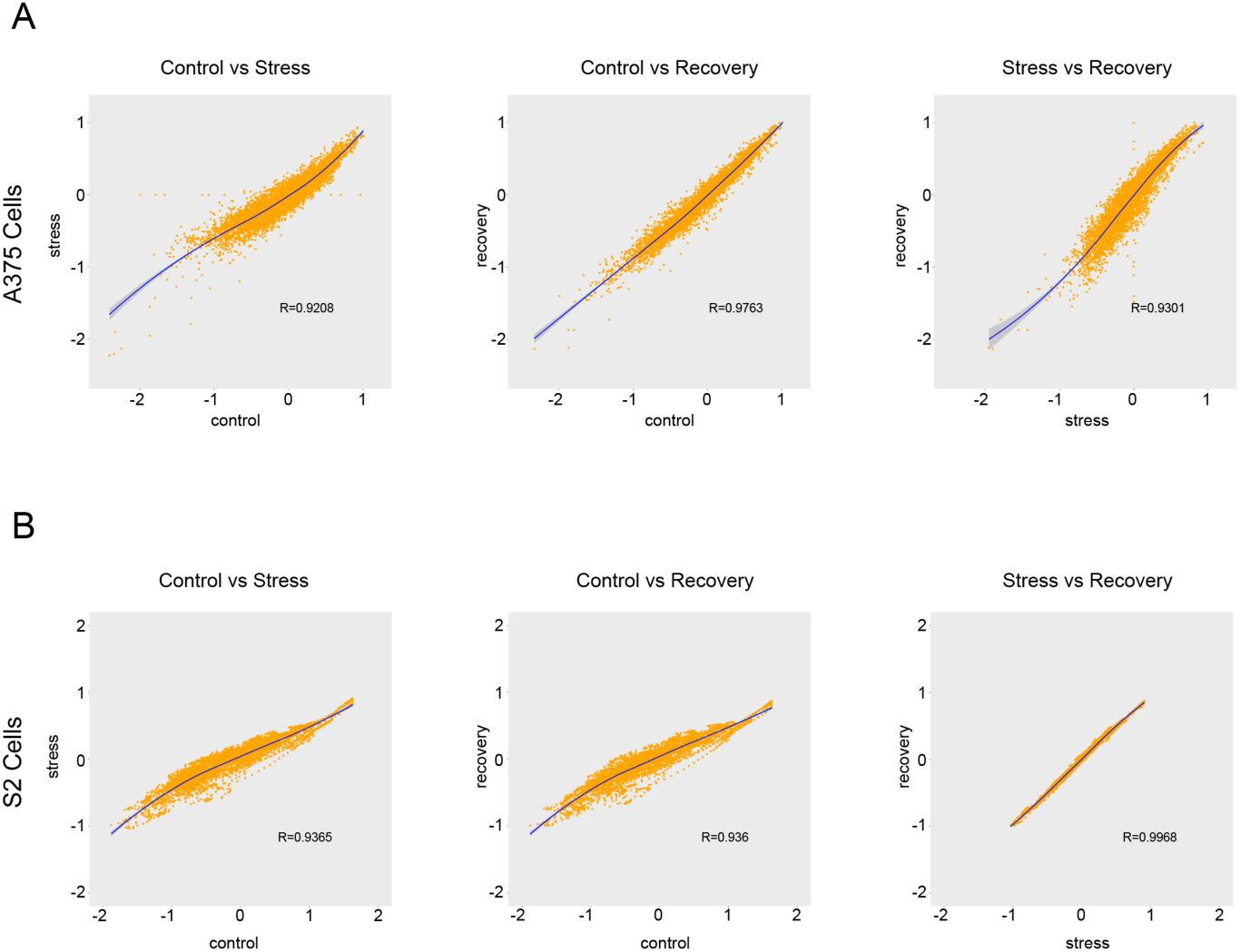
Correlation of insulation scores of control, stress and recovery in A375 and S2 cells. **A**. Correlation of control versus stress, control versus recovery, and stress versus recovery insulation scores of human A375 chromosome 1. Insulation scores were derived from 40kb matrices using the cworld-Dekker’s matrix2insulation. B. Correlation of control versus stress, control versus recovery, and stress versus recovery insulation scores of *Drosophila* S2 chromosome 2L. Insulation scores were derived from 5kb matrices using the cworld-Dekker’s matrix2insulation.

**Supplemental Figure S5:**
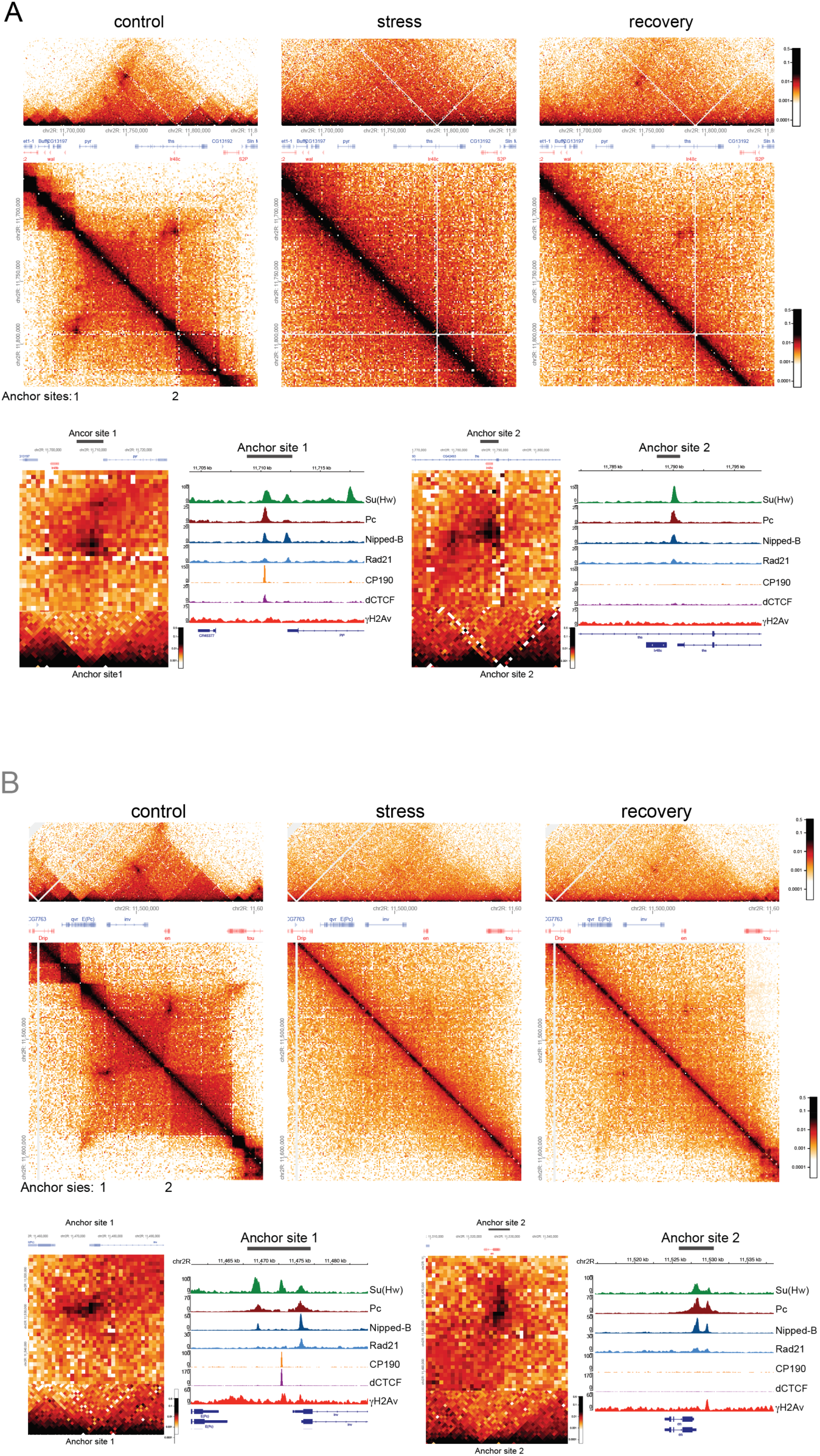
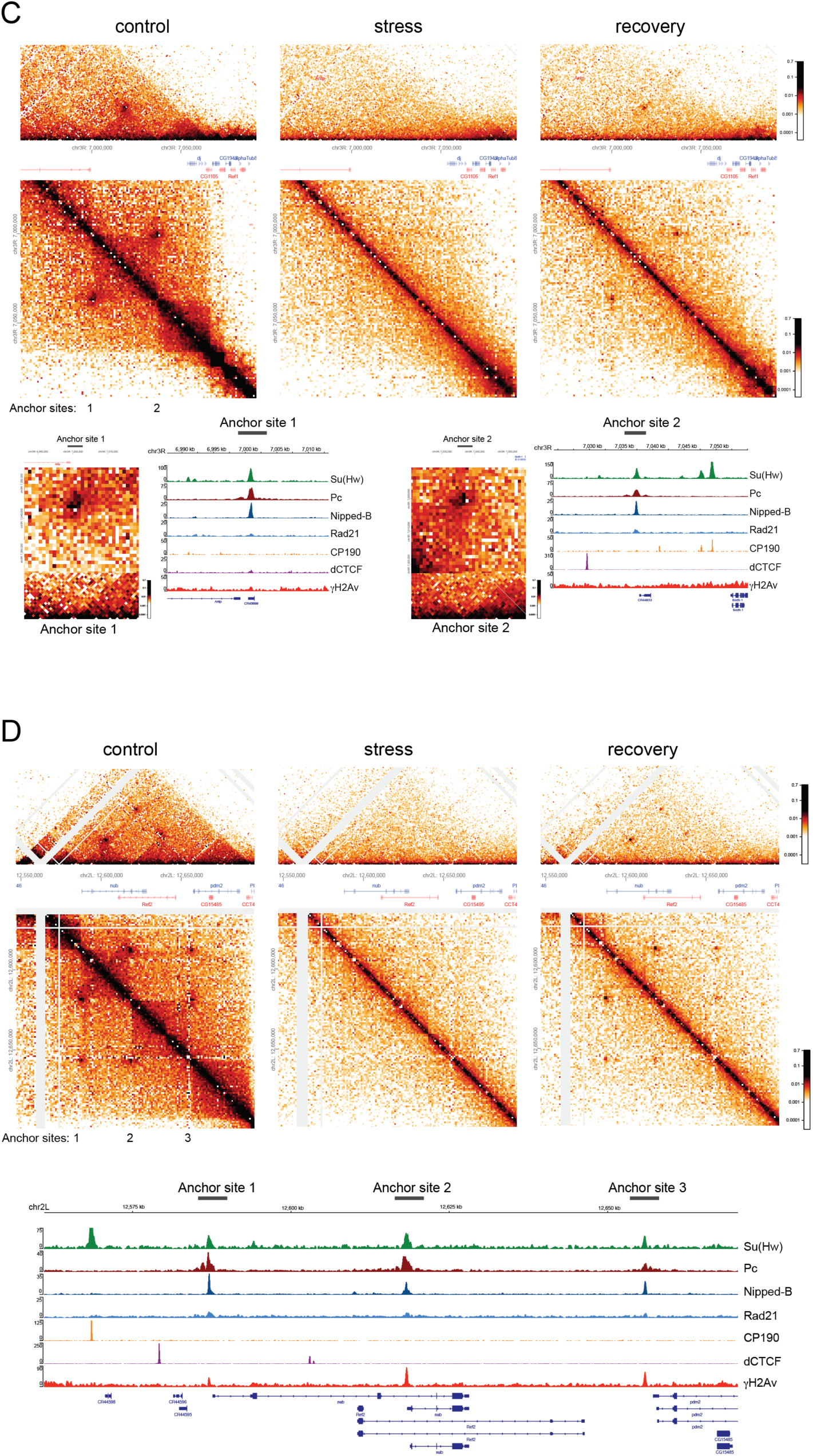
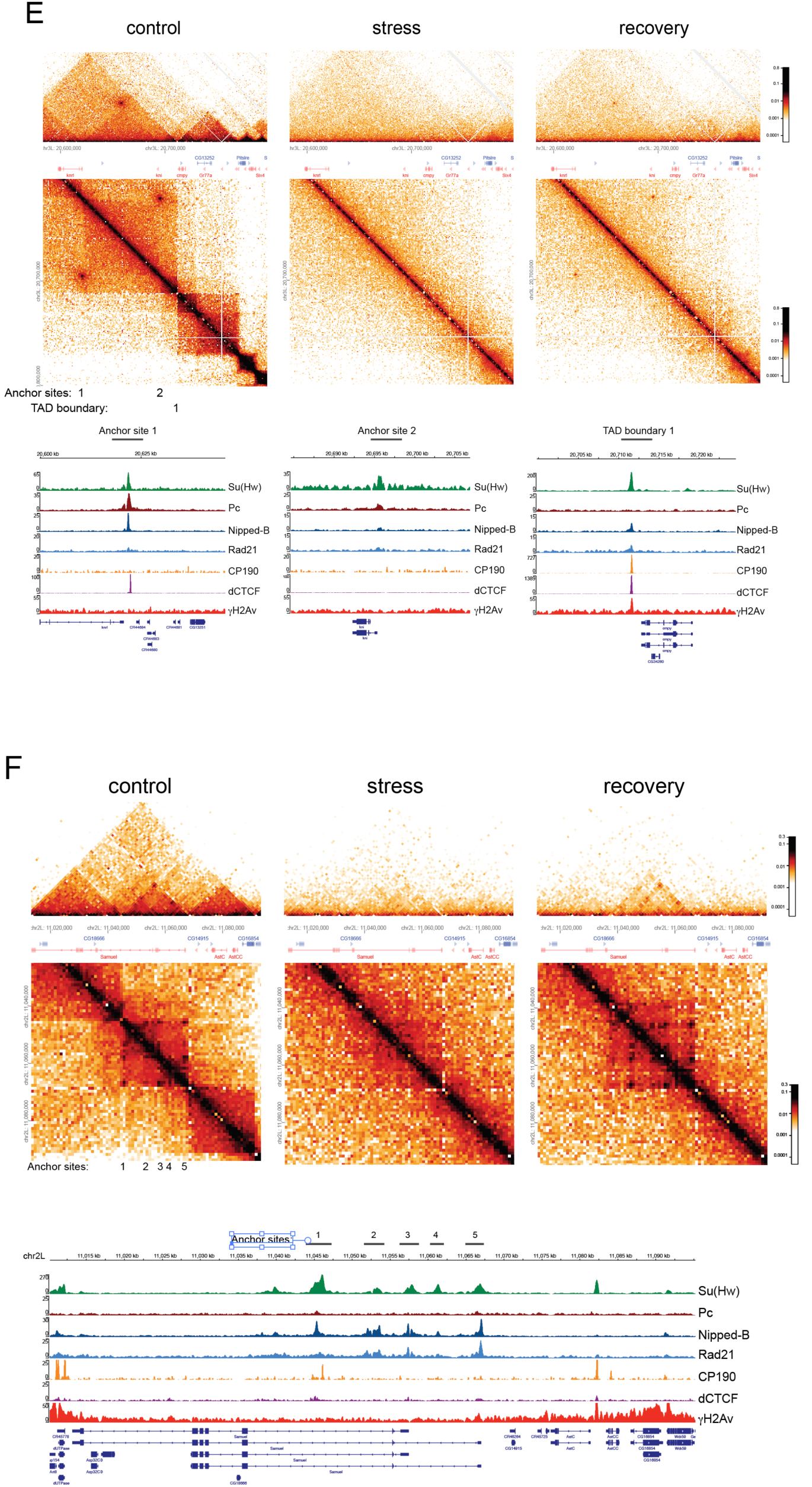
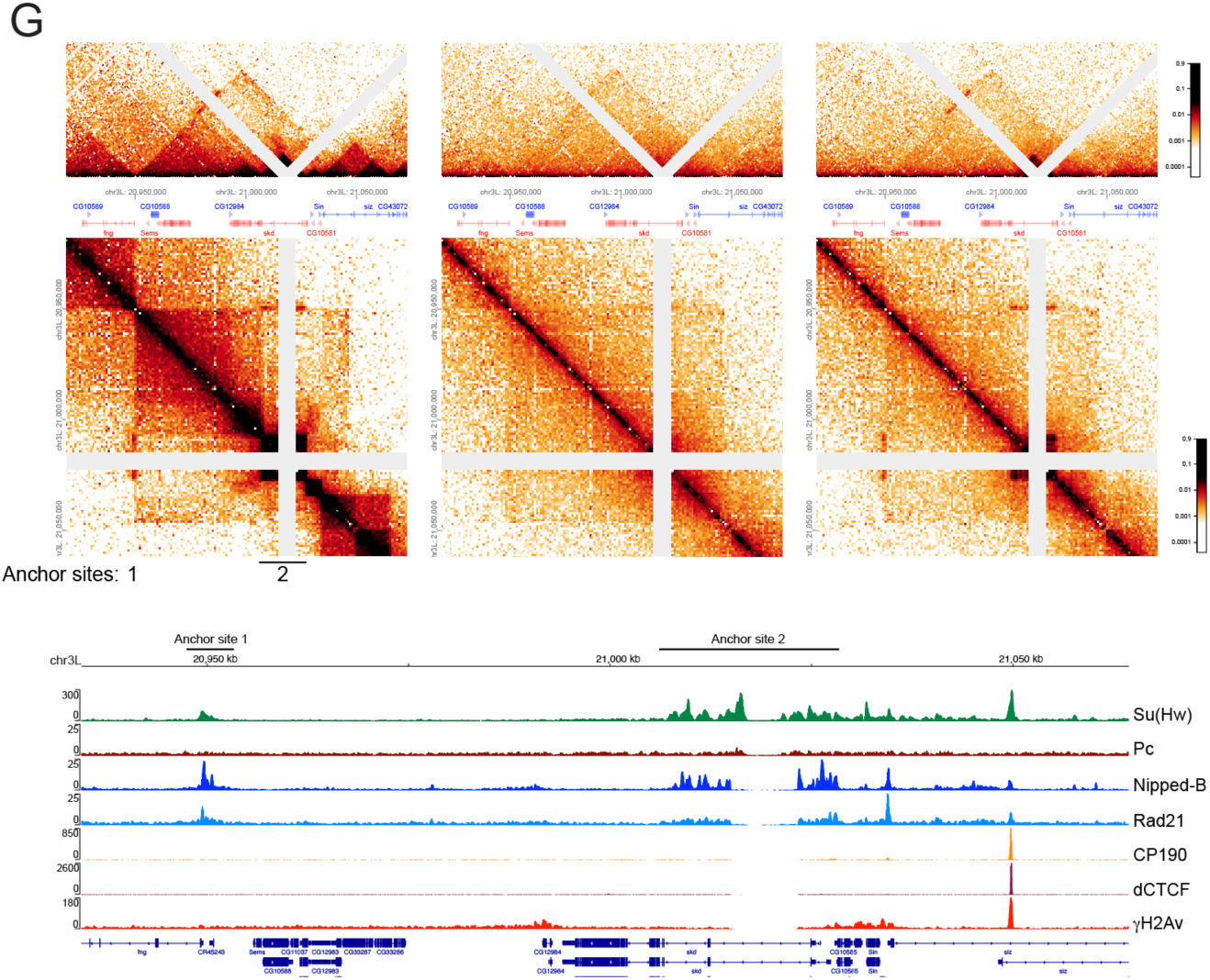
Examples of loop structures in *Drosophila*. Architectural protein ChIP-seq enrichment profiles show a common enrichment of Su(Hw) and cohesin at anchor sites. Loop structures are lost during osmotic stress and regained after recovery. (**A-E**) show enrichment of PC at anchor sites. (**F, G**) show absence of PC. (**B, D**) anchors show enrichment in γH2Av. (**A**, **B** and **C)** contact map insets show loop anchors and dots. Boundary activity is associated with anchor sites. (**E**) shows a sharp difference between loop anchors and TAD boundaries, with the TAD boundary enriched in insulator proteins cohesin and γH2Av, but not forming a loop, and not recovered after return to original media. (**G**) a continuous stretch of Su(Hw) and cohesin enrichment produces a small TAD and a stretch of long-range contacts that are dissolved during stress and reappear during recovery.

